# Gene Regulatory Programs that Specify Age-Related Differences during Thymocyte Development

**DOI:** 10.1101/2024.06.14.599011

**Authors:** Divya Ganapathi Sankaran, Hongya Zhu, Viviana I. Maymi, Isabel M. Forlastro, Ya Jiang, Nathan Laniewski, Kristin M. Scheible, Brian D. Rudd, Andrew W. Grimson

## Abstract

T cell development is fundamental to immune system establishment, yet how this development changes with age remains poorly understood. Here, we construct a transcriptional and epigenetic atlas of T cell developmental programs in neonatal and adult mice, revealing the ontogeny of divergent gene regulatory programs and their link to age-related differences in phenotype and function. Specifically, we identify a gene module that diverges with age from the earliest stages of genesis and includes programs that govern effector response and cell cycle regulation. Moreover, we reveal that neonates possess more accessible chromatin during early thymocyte development, likely establishing poised gene expression programs that manifest later in thymocyte development. Finally, we leverage this atlas, employing a CRISPR-based perturbation approach coupled with single-cell RNA sequencing as a readout to uncover a conserved transcriptional regulator, *Zbtb20,* that contributes to age-dependent differences in T cell development. Altogether, our study defines transcriptional and epigenetic programs that regulate age-specific differences in T cell development.

## INTRODUCTION

CD8+ T cells, a major component of the adaptive immune system, exhibit substantial and numerous age-dependent differences^1,2^. Neonatal CD8+ T cells, when compared to their adult counterparts, have a limited T cell receptor (TCR) repertoire, have impaired capacity to form long-lasting memory cells, possess innate-like properties, and persist into adulthood^1,3–5^. The mechanisms underlying these and other age-related differences are understood poorly. Previous studies have found that neonatal naive CD8+ T cells exhibit effector-like gene regulatory programs, which are repressed in adult cells, and these differences determine their alternative fates after activation^1^. However, when and how these age-related differences are established during CD8+ T cell development is unclear. Understanding these differences will unveil unique processes that dictate diverse CD8+ T cell developmental programs across different ages and provide insights into development-related heterogeneity within adulthood.

A fundamental difference between adult and neonatal CD8+ T cells lies in their developmental origin^1,6,7^. T cells, or thymocytes, develop in the thymus – the primary lymphoid organ – from multipotent hematopoietic stem cells (HSCs)^8–11^. HSCs arise from different anatomical niches depending on age^12–18^. In adults, bone marrow HSCs seed the thymus and differentiate into T cells^19^. Adult T-cell differentiation is a well-studied, tightly regulated, multistep developmental program that occurs in concert with the cell cycle^3,20–23^. In contrast, neonatal thymocyte development is less studied^3,24,25^. The neonatal thymus, unlike the adult thymus, not only has anatomical variations but also hosts unique HSCs that are thought to originate in the aorta-gonad-mesonephros and/or the fetal liver^26–29^. Thus, it is possible that the developmental trajectory of neonatal and adult CD8+ T cells diverges from the earliest stages of their genesis. How the distinct adult and neonatal developmental origin impinges on their respective chromatin states and generates these divergences is unknown. Alternatively, age-dependent gene regulatory programs may diverge gradually, culminating in the differences observed in mature naïve CD8+ T cells. Such age-dependent developmental programs have not been comprehensively explored.

An additional source of divergence between adult and neonatal thymocytes relates to their development. Major stages of thymocyte development in mice are demarcated by CD4, CD8, CD44 and CD25 expression, as well as by the timing of TCR gene rearrangements^30–34^. First, differentiation progresses through four CD4/CD8 double-negative (DN) stages, DN1-4, during which cells commit to T-cell lineage and rearrange their TCR genes in cell cycle arrested thymocytes^31,35–37^. Successful rearrangement relieves cell cycle arrest, and thymocytes transition to the CD4+ and CD8+ double-positive (DP) stage^38,39^. During this stage, DP thymocytes undergo positive or negative selection based on fine TCR recognition specificity, culminating in their progression to proliferative CD4 or CD8 single-positive (SP4 or SP8) T cells^40,41^. The SP8 T cells migrate into peripheral organs, such as the spleen, and complete maturation. During each of these stages, adult thymocytes undergo sequential changes in their gene expression programs that govern their developmental progression^42,43^. Presumably, many of the gene expression programs are congruent between adult and neonatal cells, as both programs generate mature CD8+ T cells; however, limited data exist to address this possibility. Moreover, there exists a stark difference in adult and neonatal thymocyte developmental rates; neonatal HSCs rapidly differentiate into mature thymocytes within 1-2 weeks, whereas their adult counterparts undergo a 3-4 week program^44–47^. How age-related gene expression programs contribute to these distinct developmental trajectories is not known.

Given the differences between adult and neonatal thymocyte origin, development and function^1,3^, we set out to compare age-related gene regulatory programs in thymocyte development. Through transcriptome profiling (RNA-seq) and chromatin accessibility profiling (ATAC-seq, assay for transposase-accessible chromatin sequencing) on adult and neonatal thymocytes at multiple stages of development, we generated a transcriptional and epigenetic atlas, linking phenotype and function. We identify an age-dependent gene module that shows consistent divergence across development and define signature genes that demonstrate age-dependent variation at distinct stages of thymocyte development. In the age-dependent gene module, genes regulating cell cycle are upregulated in neonates, while effector response genes are upregulated in adults across thymocyte development. Interestingly, the epigenetic atlas reveals that neonates have more accessible chromatin and more “poised” genes in early thymocyte development, which only manifest differential expression later during T cell maturation. We identify transcription factors whose activities likely underlie alternative gene expression programs and phenotypes in adults and neonatal thymocytes. In particular, we identify and focus on a conserved transcriptional regulator, *Zbtb20*, which is preferentially expressed in adult HSCs and during thymocyte development. We employ a CRISPR-based perturbation approach coupled with single-cell RNA sequencing as a readout to validate the role of *Zbtb20* in thymocyte development ^48,49^. In summary, our transcriptomic and epigenetic atlas reveals how age-related gene expression programs diverge during thymocyte development and defines transcriptional regulators of thymocyte development. This resource lays a foundation for dissecting mechanisms underlying age-related divergence during thymocyte development.

## RESULTS

### Age-dependent divergence in gene expression occurs early in thymocyte development

To gain insights into adult and neonate-specific T cell developmental gene regulatory programs, we isolated cells from thymi and spleens of neonatal (5-7 days post-birth) and adult (12 weeks) mice and subjected these cells to RNA-seq and ATAC-seq (Figure 1A). Antibodies to CD4, CD8, CD44 and CD25 were utilized for flow cytometry to isolate thymocytes at different stages of development^50^. We obtained cells from four stages of thymocyte development: pooled DN1-3 cells (referred to as DN hereafter), double-positive and single-positive CD8 cells (DP and SP8, respectively) and mature splenic naïve CD8+ T cells (Figures 1B and S1A). We also compared the proportions of these thymocytes and found that adults exhibited higher proportions of DN2 and DN3 thymocytes, while neonates displayed higher proportions of DP and SP8 thymocytes, suggesting age-related differences in the progression through thymocyte development^47,51,52^ (Figures 1C-1E and S1B-D). To characterize the age-related differences in gene expression, we generated four biological replicates of RNA-seq and ATAC-seq profiles from both adult and neonatal mice for all four developmental stages (DN, DP, SP8 and CD8). We first focused on the RNA-seq profiles; following routine quality control analysis, at least three biological replicates were retained for downstream analysis (Figures S2A-C).

**Figure 1.**
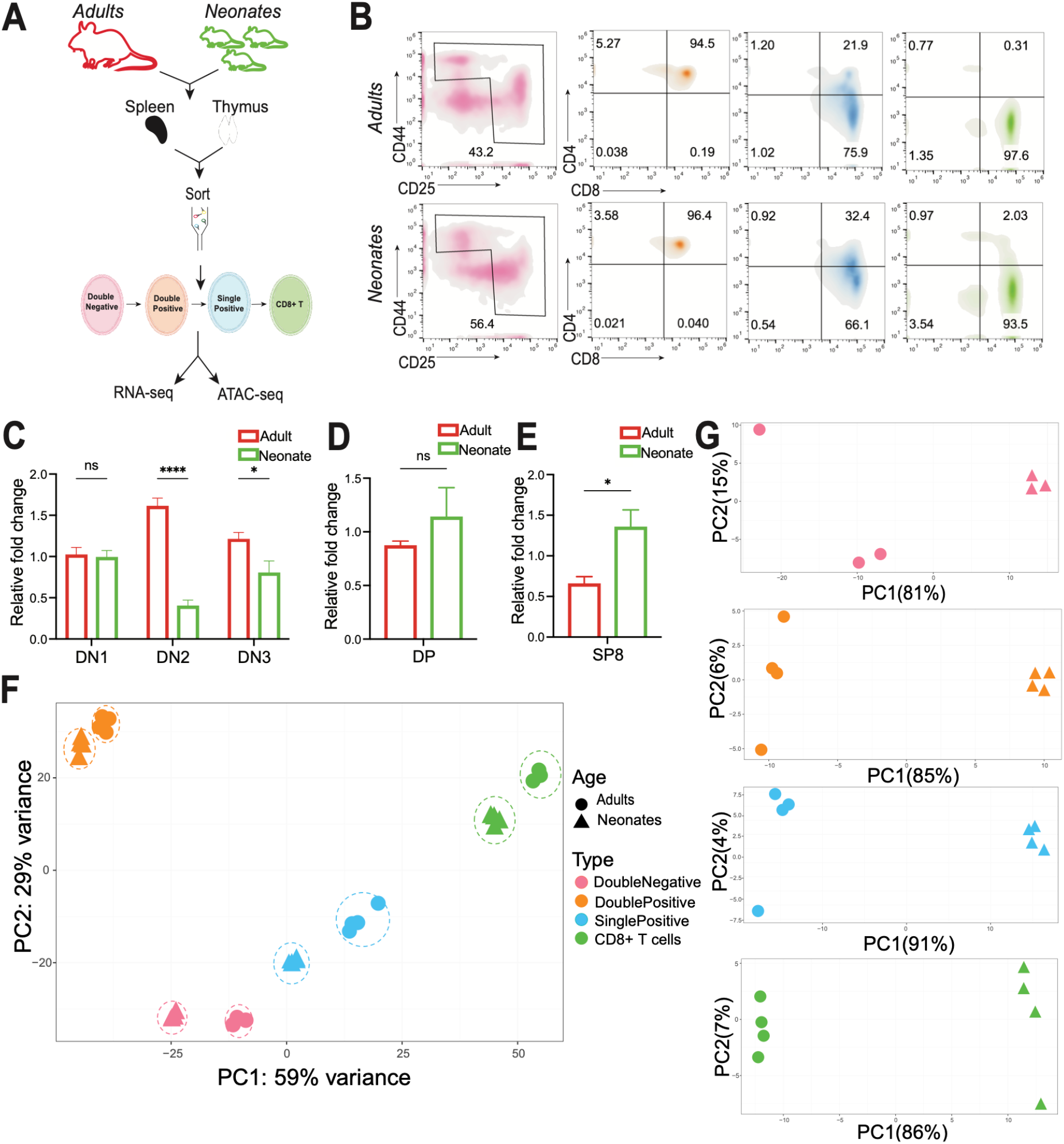
Transcriptome profiling of adult and neonatal thymocytes and splenocytes. (A) Isolation strategy of double negative (magenta), double positive (orange), single positive (blue) and CD8+ (green) thymocyte/splenocyte populations from adult and neonatal mice. (B) Flow cytometry analysis of indicated markers in adult (top) and neonatal (bottom) samples, with cell types color-coded as per panel A. (C) Relative fold change in cell numbers of adult (red bars) and neonatal (green bars) DN1, DN2 and DN3 thymocytes, compared with two-way ANOVA (ns, not significant; *, p < 0.05; **, p < 0.01, ****, p < 0.0001). (D) Relative fold change in cell numbers of adult and neonatal DP thymocytes; otherwise, as described in panel C. (E) Relative fold change in cell numbers of SP8 thymocytes post-sort; otherwise, as described in panel C. (F) Principal component analysis (PCA) of RNA-seq profiles (top 500 genes), with percentage variance associated with each axis indicated. Adult and neonatal samples denoted by circular and triangular points respectively, with developmental stage color-coded as per panel A. (G) PCA for adult (circles) and neonatal (triangles) DN, DP, SP8 and CD8+ T cell transcriptomes (panels ordered from top to bottom), with developmental stage color-coded as per panel A.

We asked when age-dependent differences in gene expression manifest. We examined gene expression programs in adult and neonatal thymocyte development and compared their transcriptome profiles using principal component analysis (PCA). Adult and neonatal naïve CD8+ T cell programs are markedly different^4,53^, suggesting that perhaps the major variance between thymocyte transcriptomes would arise due to animal’s age. Instead, the PCA of RNA-seq profiles revealed that thymocytes cluster primarily based on developmental stage, although age also contributed to the total variation (2.6%; Figure S2D). In addition, biological replicates at each stage correlated strongly (Pearson coefficient, *r* > 0.975; Figures 1F and S2E). Nevertheless, when examining individual stages, samples segregated strongly (>80% of the variance) by age (Figures 1F and 1G). These observations suggest that thymocyte development entails gene expression changes that occur in concert for adults and neonates, with additional age-specific differences overlayed. Indeed, age-specific expression programs are evident at the DN stage, the earliest stage we examined, and across the complete developmental trajectory (Figure 1G).

### Thymocyte gene expression programs determined by developmental origin

Given that age-specific expression programs are evident early and at every developmental stage, we asked which genes show consistent age-dependent expression differences across thymocyte development. Such a gene set may reveal how developmental origin impinges on age-dependent gene regulatory programs during T cell development. We first identified significant differentially expressed genes (DEGs) between adults and neonates at each of the four developmental stages and then intersected them to define this set (Figure 2A and S3A). Comparing DEGs between adults and neonates at each stage, the DN and the SP8 stages show the highest number of DEGs, while the DP has the lowest. This observation is consistent with the broad shutdown of several transcriptional programs that occurs during the DP stage (Figure 2A and S3A)^42,54^. We then intersected DEGs from all four stages, yielding a set of 275 genes that have consistent age-related differences throughout thymocyte development (Figures 2A, 2B, S3A and Table S1), which we refer to as the age-dependent gene module.

**Figure 2.**
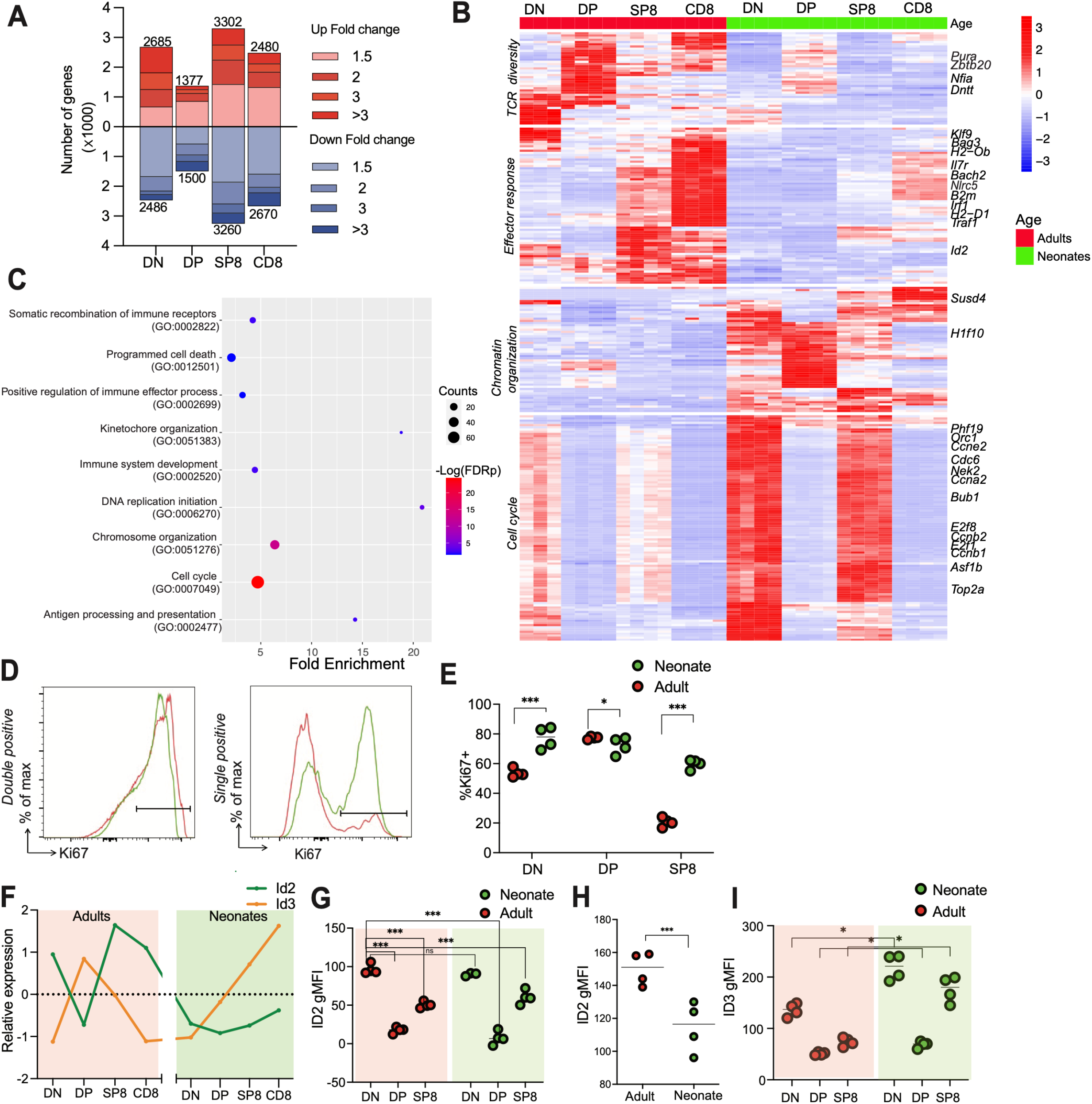
Gene regulatory programs in adult and neonatal thymocytes. (A) Number of significant DEGs (y-axis) between adults (red) and neonates (blue) at each developmental stage (x-axis; Wald test, FDR adjusted p <0.05). (B) Heatmap of age-dependent gene module, comparing expression (row normalized z-scores; see color-coded scale bar) in adults (left) and neonates (right). (C) Significant GO biological processes in age-dependent gene module, analyzed by PANTHER overrepresentation test (Fisher’s exact test, FDR adjusted p <0.05), focusing on repetitive and relevant GO biological processes. (D) (Left) Ki67-positive population in adult (red) and neonatal (green) DP thymocytes. (Right) Ki67-positive population in adult (red) and neonatal (green) SP8 thymocytes. (E) Percentage of Ki67-positive population in adult (red) and neonatal (green) thymocytes. Two-way ANOVA test (*, p < 0.05, ***, p < 0.001). (F) Id2 (green) and Id3 (orange) normalized transcript expression (z-scores) in adult (left panel) and neonatal (right panel) thymocyte development. (G) Id2 geometric Mean Fluorescence Intensity (gMFI) in adult (red circles) and neonatal (green circles) thymocytes, compared with two-way ANOVA test (ns, not significant; *, p<0.05, ***, p<0.0001). (H) Id2 gMFI in adult and neonatal DN1 thymocytes; otherwise, as described in panel G. (I) Id3 gMFI in adult and neonatal thymocytes; otherwise, as described in panel G.

To characterize the age-dependent gene module, we performed hierarchical clustering (across all samples) and functional annotation on these genes (Figures 2B, 2C, S3B and Table S2). The largest cluster, comprising 117 genes, exhibits higher expression in neonates than in adults (Figure 2B; bottom). Intriguingly, in both adults and neonates, this cluster shows a distinct pattern: high expression in DN thymocytes, which reduces in DP thymocytes and then rises back to high levels in SP8 thymocytes before reaching the lowest levels in CD8+ T cells. These dynamics are more pronounced in neonates (Figure 2B). Strikingly, most of the genes in this cluster are established cell cycle genes (75 of 117 genes), including *Ccne2*, *Top2a*, *Mcm10*, *Orc1*, *Ccna2*, *Ccnb2*, *Cdc6*, *Bub1 and Nek2* (Figures 2B, 2C and S3C)^55–61^. A two-fold change in many of these genes is sufficient to modulate the cell cycle^61–67^. These results indicate that neonatal thymocytes exhibit differential control of cell cycle genes, an observation that suggests enhanced proliferation during DN and/or SP8 stages of thymocyte development.

To investigate if neonatal thymocytes are more proliferative than adults, we used flow cytometry to examine the expression of Ki67, a cell proliferation marker, in the adult and neonatal thymocytes^68^. We found an increased proportion of Ki67-positive cells in neonatal DN and SP8 stages (Figures 2D and 2E). Neonates have slightly lower proportions of Ki67-positive cells than adults during the DP stage, consistent with the cell cycle genes’ expression being least divergent in DP thymocytes (Figures 2B, 2D and 2E). These findings suggest that neonates are more proliferative than adults at DN and SP8 stages of thymocyte development^23,69–71^. Additionally, amongst the genes preferentially expressed in neonates, we identified genes relevant to chromatin organization, including *H1f10* and *Asf1b* (Figures 2B and 2C)^72,73^. These observations suggest that neonates preferentially express genes that regulate chromatin organization and cell cycle regulation across thymocyte development.

The age-related module also contains 44 genes that are upregulated in adults compared to neonates (Figures 2B; top). Prominent amongst these is a cluster containing *Il7r*, *Id2*, *Bach2 and Irf1*, which are essential for thymocyte development and effector response (Figures 2B and 2C). For example, *Id2,* a co-transcriptional inhibitor of E-proteins that regulates thymocyte development, effector response and cell cycle^74^, exhibits preferential expression in adults, dovetailing with the downregulation of E-proteins *E2f1* and *E2f8* in adults (Figures 2B and 2C). Interestingly, *Id2* expression in adults contrasts with that of *Id3,* another Id-family transcription factor, which acts antagonistically to *Id2*^74–77^. *Id2* is higher in adults and *Id3* is higher in neonates (Figures 2B, 2F, S3D and S3E). We next examined if the expression differences correlate with protein levels using flow cytometry. Id2 expression is higher in adult DN thymocytes, particularly in the DN1 stage, relative to neonates and other stages of thymocyte development (Figure 2G and 2H). In contrast, Id3 protein levels are higher in neonates than in adults (Figure 2I). These results indicate that Id2 and Id3 show age-dependent differential expression and may contribute to the gene regulatory differences between adult and neonatal thymocytes. In addition to the effector response cluster, we found that *Dntt*, which is essential for generating TCR diversity, has lower expression during neonatal thymocyte development, consistent with the reduced TCR diversity typical of neonatal T cells (Figures 2B and 2C)^73–76^.

Finally, 24 transcription factors are found within the age-dependent gene module (referred to as age-dependent TFs), including some with no known roles in age-related differences in thymocyte development. This set includes *Id2*, *Bach2*, *Rxra*, *E2f1*, *E2f8*, *Hoxa*, *Zbtb20*, *Zbtb42, Pura*, *Nfia*, *Klf9* and *Klf11.* Amongst these TFs, *Id2, Bach2, E2f family* and *Hoxa* have well-studied roles in adult T cell development^74,77–83^. In contrast, *Zbtb20*, *Zbtb42*, *Pura*, *Klf9* and *Klf11* have no established roles in thymocyte development. Interestingly, *Zbtb20* is known to regulate the adult effector response and immunological memory formation in mature CD8+ T cells ^84^. Whether and how these TFs regulate age-related differences in thymocyte development is unclear. Overall, our analyses show that adults and neonates exhibit consistent age-dependent divergence in transcriptional regulation of cell cycle control and effector response. These programs, likely regulated by *Id* genes and/or other age-dependent TFs, differ from the earliest thymocyte developmental stages, reflecting programs that likely derive from their distinct developmental origins.

### Adult and neonatal thymocyte stage-specific gene expression programs

We next sought to identify genes whose expression differs between adults and neonates only at specific thymocyte developmental stages. Such gene sets may reveal age-dependent divergences during individual developmental stages and also highlight temporal regulation of gene regulatory programs. First, we performed differential expression analysis, comparing all adult and neonatal samples while using developmental stages as an additional variable in the analysis. We identified 3,369 differentially expressed genes, with 1,978 upregulated in adults and 1,391 in neonates (Figure S4A); hierarchical clustering allowed us to identify gene sets upregulated in either adults or neonates at each developmental time point (Figure S4A).

We first considered gene sets upregulated at individual adult developmental stages (Figure 3A and Table S3). Analyzing the adult upregulated genes using functional annotation revealed a strong enrichment for immunity-related gene sets, including T-cell differentiation and T-cell function, implying variations in these processes between adults and neonates (Figures 3C, S5A and Table S4). These differences emerge as early as the DN stage and are linked to stage-specific processes at each stage. For instance, during the DN stage, repression of progenitor (*Flt3*, *Spi1*, *Nfam1*, *Rgs1* and *Bcl11a)* and non-T cell lineage (*Id2*, *Mef2c*, *Bank1, Tyrobp and Ctla4)* genes occurs. These genes are included in the adult DN (ADN) signature gene set and have reduced expression in the neonatal DN stage (Figures 3A; ADN cluster, 3C and S5B) suggesting that neonates may control the shutdown of these progenitor programs and non-T cell lineage programs earlier than adults, consistent with previous results, and possibly due to their distinct developmental origin^2,85–90^. Likewise, during the DP stage, thymocytes undergo positive selection or apoptosis based on TCR interaction affinity and migrate to the thymic medulla to become SP8 cells^91,92^. Genes that regulate apoptotic and migratory processes, including *Nfat5*, *Egr2*, *Tec*, *Pdk1*, *Bik*, *Eya2*, *Zbtb20*, *Cxcr4*, *Cnn3*, *Cldn10, Tead2* and *Cacnb3*, are upregulated in the adult DP stage (ADP) and are reduced in neonatal DP stage, suggesting possible age-related alterations to selection^93^ and migration (Figures 3A; ADP cluster and 3C)^94–97^.

**Figure 3.**
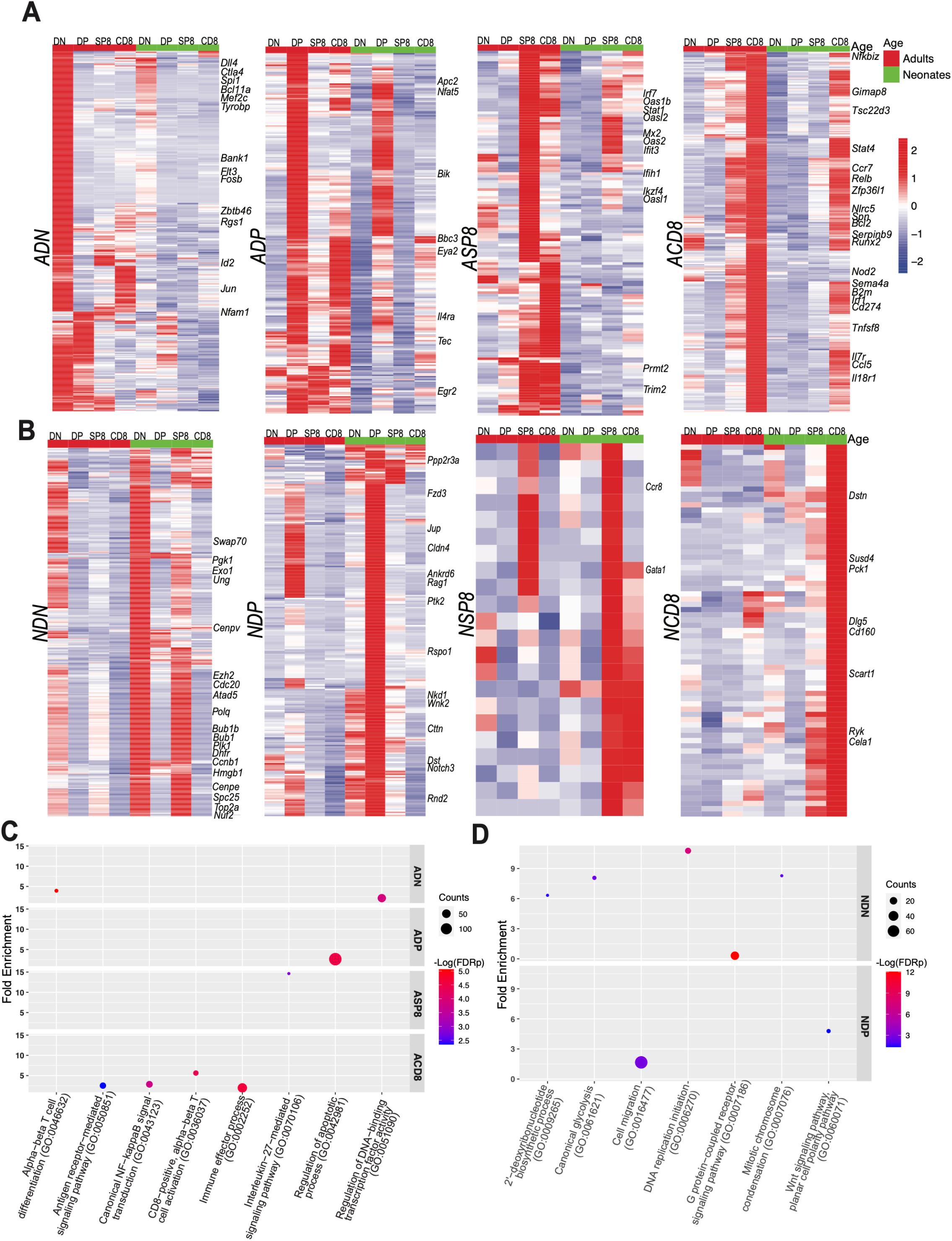
Adult and neonatal thymocyte stage-specific gene expression programs. (A) Heatmap of adult stage-specific signature genes, comparing expression (row normalized z-scores; see color-coded scale bar) in adults (left) and neonates (right). (B) Heatmap of neonatal stage-specific signature genes, otherwise, as described in panel A. (C) Significant GO biological processes in adult signature genes, analyzed by PANTHER overrepresentation test (Fisher’s exact test, FDR adjusted p<0.05), focusing on repetitive and relevant GO biological processes. (D) Significant GO biological processes in neonatal signature genes, otherwise, as described in panel C.

Additional differences in T cell function between adults and neonates were evident through examination of the adult CD8 (ACD8) signature gene set. Genes in the ACD8 cluster include those that regulate T cell activation, TCR signaling and effector response (*Il7r*, *Runx2*, *Bach2*, *B2m, Cd69, Stat4*, *Zfp36l1*, *Junb*, *Il2rb*, *Spn* and *Rora*)^98,99^. These genes have reduced expression in neonatal CD8+ T cells (Figures 3A; ACD8 cluster and 3C). *NF-κB* signaling pathway genes (*Nfkbiz*, *Rel*, *Relb*, *Tnf, Bcl2*, *Traf6* and *Tnfsf10)* were also downregulated in neonatal CD8+ T cells, relative to adult counterparts (Figures 3A; ACD8 cluster and 3C)^100–103^. Differences in these programs may specify age-related differences in T cell function, including activation and effector response. Together, these findings highlight upregulated programs in adults relative to neonates.

We next analyzed gene sets upregulated at individual neonatal developmental stages (Figures 3B and Table S5), revealing a significant enrichment in cell cycle-related processes and cell migration during neonatal thymocyte development (Figures 3B, 3D, S5C and Table S6). In the neonatal DN (NDN) signature gene set, we observe two distinct clusters: one with high expression at the DN stage and a second with high expression at both the DN and SP8 stages. Multiple genes essential to cell cycle progression, including *E2f1*, *E2f3*, *E2f8*, *Foxm1, Ccnb1*, *Cenpe*, *Plk1*, *Top2a* and *Cdc25a*, show high expression in the latter cluster (Figures 3B; NDN cluster)^55–60^. Additionally, the neonatal DN signature gene set includes genes that regulate metabolic processes (*Rxra* and *Pparg*), including glycolysis (*Pgk1*, *Gapdh*, *Pkm*, *Grhpr*, *Dhfr* and *Tpi1*) ^104,105^, suggesting that neonatal thymocytes are not only more proliferative but also exhibit altered metabolism during the DN stage (Figures 3B; NDN cluster and 3D)^106^.

During the DP stage, the *Wnt* signaling pathway, known to regulate cell migration and cell division, shows elevated expression in neonatal DP (NDP) thymocytes, implicating disparities in *Wnt* signaling-mediated migration and cell division between neonates and adults (Figures 3B; NDP cluster and 3D) ^94–96^. In the neonatal SP8 (NSP8) and CD8 (NCD8) signature gene sets, fewer genes including *Gata1*, *Ccr8*, *Susd4* and *Cd160* have established roles in T cell biology, while others, such as *Scart1*, *Dstn*, *Ryk* and *Cela1* do not have known roles in neonatal thymocyte development (Figures 3B; NSP8, NCD8 clusters and 3D).

Collectively, adults and neonates differ in the expression of genes that regulate T cell commitment and T cell-cycle related processes in the DN stage, TCR-based selection and T cell migration in the DP stage and T cell function, later in thymocyte development. These distinctions arise as early as the DN stage and are linked to stage-specific T-cell developmental pathways. For many of these pathways, the related genes are upregulated in adults. In neonates, we find a global elevation of cell cycle-related genes. How these adult and neonatal signature genes are temporally regulated to achieve developmental stage-and age-specificity is not known.

### Adult and neonatal chromatin landscapes across thymocyte development

To better understand stage-specific gene expression during adult and neonatal thymocyte development, we performed ATAC-seq on the same samples profiled by RNA-seq (Figures S6A-D). First, we examined the global relationships between accessible chromatin regions in adult and neonatal thymocytes. PCA of ATAC-seq profiles displayed clustering based on the developmental stage (Figure 4A), with biological replicate profiles exhibiting strong concordance (Figures 4A, S6A and S6B). Furthermore, consistent with the transcriptome data (Figure 1F), the ATAC-seq profiles of adults and neonates are clearly distinct at each developmental stage (Figures S6C and S6D).

**Figure 4.**
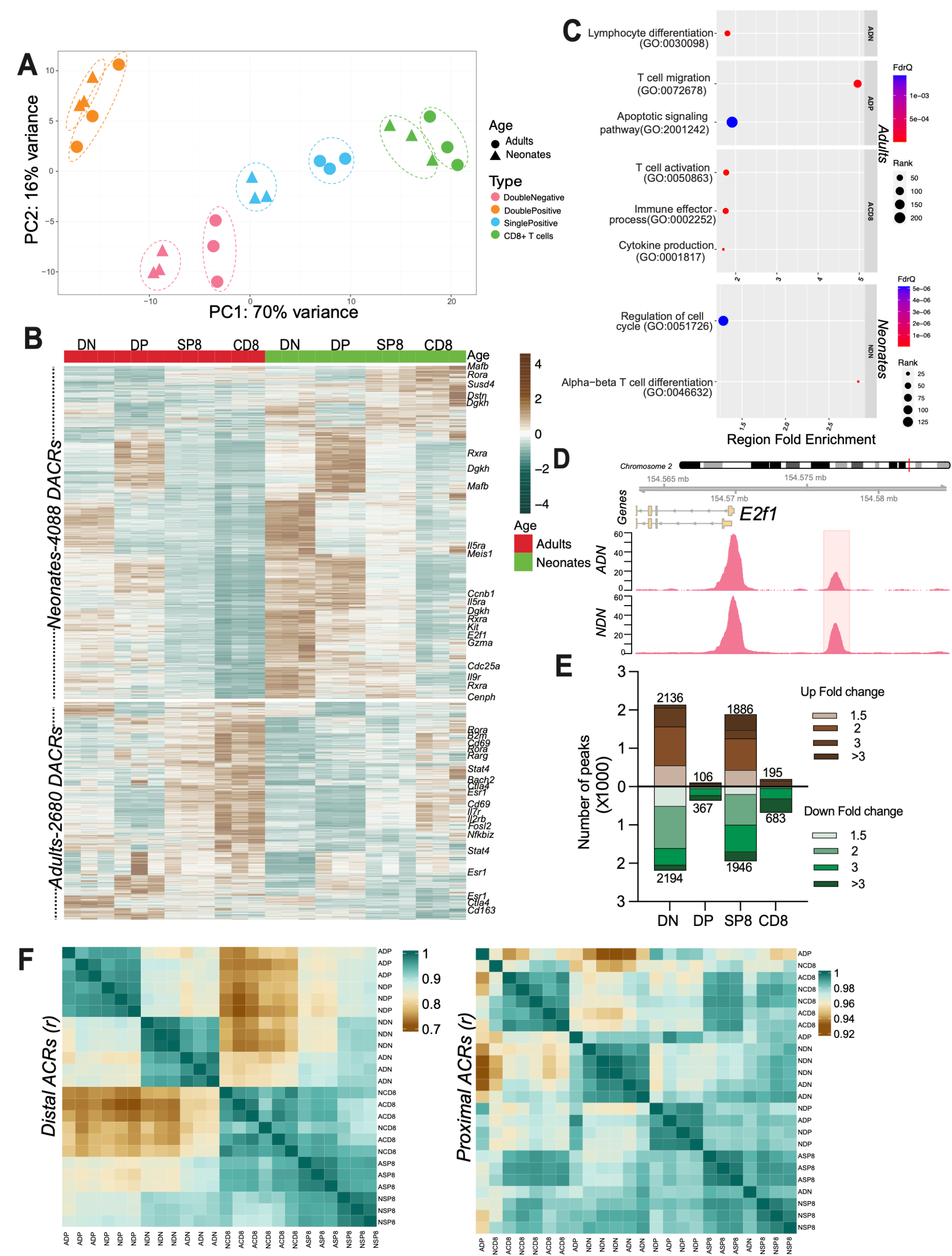
Chromatin accessibility during thymocyte development. (A) Principal component analysis (PCA) of ATAC-seq profiles (top 500 genes), with percentage variance associated with each axis indicated. Adult and neonatal samples are denoted by circular and triangular points, respectively, with developmental stage color-coded, as described in Figure 1A. (B) Heatmap of significant differential ACRs (DACRs) between adult (left) and neonatal (right) thymocytes; see color-coded scale bar, labeling differentially accessible genes associated with ACRs with correlated stage-and age-specific expression changes (Wald test, FDR adjusted p <0.05). (C) Significant GO biological processes in adult and neonatal DACRs, analyzed by GREAT binomial test (FDR adjusted p <0.05), focusing on processes that explain temporal regulation in signature genes. (D) ATAC-seq signal tracks of the indicated gene (E2f1) in adult and neonatal DN cells. Tracks are group-scaled per dataset. (E) Number of significant DACRs between adults (brown) and neonates (green) at different thymocyte developmental stages (x-axis; Wald test, FDR adjusted p <0.05). (F) (Left) Pearson correlation (r) between distal ACRs in adult and neonatal thymocytes (see color-coded scale bar). (Right) Pearson correlation (r) between proximal ACRs in adult and neonatal thymocytes (see color-coded scale bar).

To compare the accessibility differences between adults and neonates, we performed differential accessibility analysis between adults and neonates across all thymocyte samples, using the developmental stage as an additional variable in the analysis (Figure 4B). Our data indicates that neonates have more accessible chromatin regions (ACRs) in terms of magnitude and number during early thymocyte development, while adults have highly accessible regions in late thymocyte development (Figure 4B). These results suggest that adult and neonatal thymocytes harbor distinct chromatin accessibility landscapes, with neonates having more accessible chromatin during early thymocyte development, possibly due to their different developmental origins.

We next asked which age-and stage-specific gene expression programs show congruent changes in chromatin accessibility. Functional annotation analysis revealed congruent shifts in processes including thymocyte differentiation in the adult DN stage (*Id2*, *Ctla4, Klf9* and *Cd163)*^74,77,107–110^, thymocyte migration and apoptosis in the adult DP stage (*Pdk1*, *Cxcr4*, *Cnn3*, *Cldn10*, *Lag3* and *Cacnb3*)^111,112^, and activation, effector response and cytokine production in adult CD8+ T cells (*Bach2*, *Il7r, Bcl2, B2m, Il2rb, Rora, Lta, Cd69, Cd96, Cd226, Stat4, Slamf7, Spn*, *Rarg* and *Zfp36l1)*^82,113–116^. The increased expression of these genes in adults relative to neonates is associated with increased accessibility (Figures 4B, 4C and 3A). Notably, in neonates, cell cycle genes including *E2f1, Ccnb1, Cdk1, Cdk2, Cdc6, Aurkb, Cenph* and *Ccdc25* are more accessible in the DN stage, congruent with the increased expression of cell cycle genes, suggesting a role in neonatal cell cycle transcriptional regulation and proliferation (Figures 4B, 4C, 4D, 2B, 3B and S6E). Moreover, neonatal thymocytes show increased accessibility around genes such as *Kit, Rxra* and *Ptcra*, possibly driving their early developmental expression. Finally, in neonatal CD8+ T cells, *Susd4*, *Dstn* and *Dgkh* show increased expression and accessibility (Figure 4B and 3B). These findings suggest that changes in chromatin accessibility contribute to age-and stage-specific gene expression. Furthermore, when we analyzed differential ACRs (DACRs) between individual developmental stages, we observed that adults and neonates show a reduced number of DACRs at DP and CD8 stages relative to DN and SP8 stages (Figure 4E). Strikingly, this reduction in DACRs during the DP stage correlates with the reduction in DEGs and transcriptional shutdown observed in the DP stage, suggesting that chromatin landscape alterations may underlie the transcriptional shutdown in the DP stage (Figures 4E and 2A).

The differential accessibility between adults and neonates also prompted us to investigate whether peaks proximal to transcriptional start sites (TSS; promoter regions) or distal (enhancers) peaks contribute to differential accessibility between adults and neonates (Figure 4F). Age-dependent changes at distal regions would likely require changes in the cognate TFs binding accessible regions. Clustering based on distal (> 2kb from the nearest TSS) and proximal ACRs (< 2kb from the nearest TSS) revealed that distal ACRs contribute significantly to the distinction between developmental stages and age, particularly evident during DN and SP8 stages; in contrast, adult and neonatal proximal ACRs were highly correlated and contributed less to the distinction (Pearson coefficient distal *r* > 0.7, proximal *r* > 0.9, Figure 4F). Hence, distal peaks play a more prominent role in distinguishing age-and stage-related transcriptional regulation than proximal TSS peaks. Collectively, our findings demonstrate that the chromatin landscape differs between adults and neonates, with neonates exhibiting more accessible chromatin during early thymocyte development. Moreover, there exists a temporal correlation between specific gene expression programs and their chromatin accessibility landscape at distinct developmental stages and ages.

### Age-related poised programs during thymocyte development

Given the high accessibility in neonatal chromatin in the DN stage, we asked whether this corresponds to an increase in DEGs (Differentially Expressed Genes) at this stage. Surprisingly, there is not a corresponding increase in neonatal DEGs during the DN stage (Figure 4B, 4C and 2A). Moreover, while certain gene expression programs correlated with their accessibility profiles, the mechanism underlying the upregulation of other stage-specific programs remained unclear (Figure 4B, 4C, 4E and 2A). We hypothesized that neonates might harbor genes that are maintained in a "poised" state and are expressed later in thymocyte development. These genes maintain an open chromatin state and are not differentially expressed in early development but become so in mid-or late-thymocyte development, potentially enabling precise temporal regulation of gene expression programs. To test this hypothesis, we defined three developmentally relevant transitions in adults and neonates: early-thymocyte development, assessed by comparing DN versus DP programs, mid-thymocyte development, comparing DP versus SP8, and late-thymocyte development, comparing SP8 versus CD8 (Figure 5A). We then identified genes that were differentially accessible but not differentially expressed during early-thymocyte development and overlapped this set with genes differentially expressed in later stages (mid-and late-thymocyte development). We refer to genes as “poised” when they are differentially accessible at a stage preceding their differential expression.

**Figure 5.**
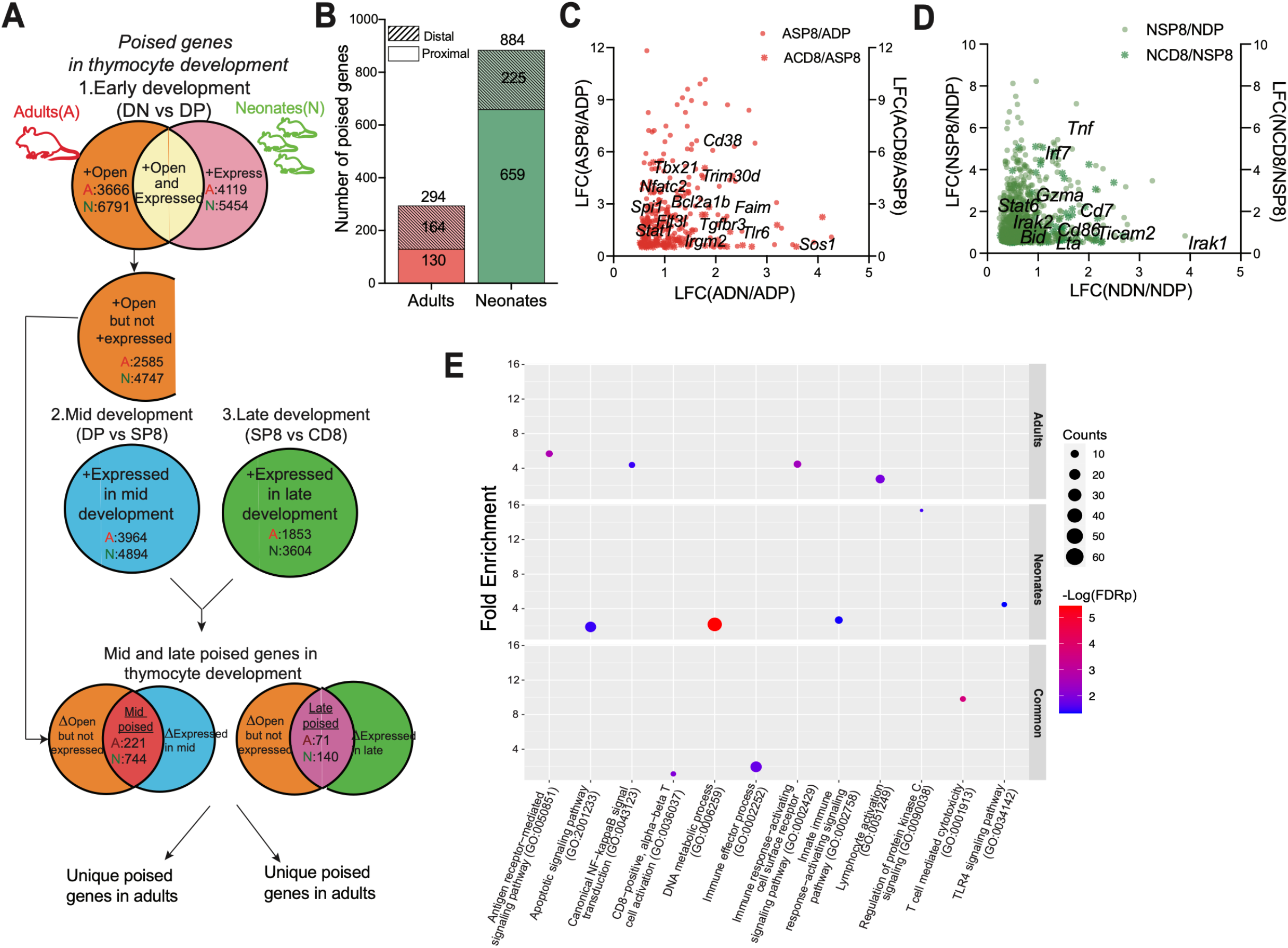
Poised genes in adults and neonates during thymocyte development. (A) Schematic of “poised genes” classification during thymocyte development. (B) Number of poised genes in adults and neonates during thymocyte development. (C) Relative expression changes and accessibility fold changes (y-and x-axis, respectively; log2 of fold change; LFC) of unique adult poised genes. (D) Relative expression changes of unique neonate poised genes; otherwise, as described in panel C. (E) Significant GO biological processes in adult and neonatal poised genes, analyzed by PANTHER overrepresentation test (Fisher’s exact test, FDR adjusted p<0.05), focusing on processes that explain temporal regulation in signature genes.

There are twice as many differentially accessible ACRs (DACRs) in neonates as compared to adults during early-thymocyte development (6,791 for neonates vs. 3,666 for adults, Figure 5A). However, the number of DEGs in adults and neonates in this early stage is similar (5,454 for neonates vs. 4,119 for adults; 1.3-fold, Figure 5A). Consequently, many more genes in neonates are accessible early relative to adults but not differentially expressed. Conversely, in late-thymocyte development, neonates have many more DEGs than adults (3,604 vs. 1,853; 2-fold, Figure 5A). We identified poised genes by intersecting DEGs in mid-and late-thymocyte development with the genes that are differentially accessible but not expressed in early-thymocyte development (Figure 5A). Consistent with our hypothesis, neonates harbor 884 poised genes (659 proximal ACRs and 225 distal ACRs, Figure 5B), compared to only 294 unique poised genes in adults (130 proximal ACRs and 164 distal ACRs, Figure 5B). This suggests that poising plays a relatively larger role in facilitating temporal control of transcriptional programs in neonates than in adults.

Functional annotation analysis revealed distinct pathways enriched in adult and neonate unique poised genes. Adult poised genes encompassed T cell activation (*Tbx21*, *Nfatc2*, *Spi1*, *Cd38* and *Flt3l), NF-κB* signaling (*Fasl*, *Stat1* and *Faim*), and antigen-receptor mediated signaling (*Themis2*, *Sos1* and *Klri2*; Figures 5C and 5E), suggesting their involvement in temporal control of *NF-κB* signaling and T cell activation in adult CD8+ T cells, dovetailing with them being ACD8 signature genes (Figure 3A; ACD8 cluster). Conversely, neonate-poised genes included innate immune response (*Irf7, Irak2, Cd86, Ticam2* and *Tyrobp*), apoptosis regulation (*Bid*, *Tnf*, *Casp8ap2*), and metabolism-related genes (Figures 5D and 5E). These findings imply that the poising of distinct adult and neonate genes may regulate age-dependent gene regulatory programs. Moreover, we also identify shared poised genes between adults and neonates, which includes immune effector processes and T cell cytotoxicity genes, indicating that effector processes are poised regardless of age (Figure 5E). Overall, we find that neonates not only have more poised genes than adults but also have distinct functions associated with poised genes.

### Functional validation of transcription factors controlling age-related differences in T cell developmental rates

Given the age-related differences in chromatin accessibility, we explored TFs that exhibit differential binding between adults and neonates at all four developmental stages, and that may, therefore, be involved in generating the age-related differences in gene expression programs. Using diffTF, which integrates chromatin accessibility and expression data, we assessed changes in TF binding activities (Figures 6A and S7A-D) ^117^ ^118^. Key developmental TFs such as *Gata3*, *Tcf7, Spi1* and *Runx3* show an altered association with the accessible regions in adults and neonates, including but not limited to the DN stage, suggesting distinct control of early T cell commitment genes^35,36,118,119^. TFs that regulate global chromatin architecture, such as *Ctcf*, show differential binding between adults and neonates across thymocyte development, suggesting a possible role for *Ctcf* in age-related chromatin landscape differences^120^. Additionally, several age-dependent TFs from the age-dependent gene module, which have no known roles in age-related differences during T cell development, such as *Zbtb42*, *Plag1, Nfia* and *Irf1,* are predicted to be differentially active when compared between adult and neonatal thymocytes (Figure 6A and Figure 2B).

**Figure 6.**
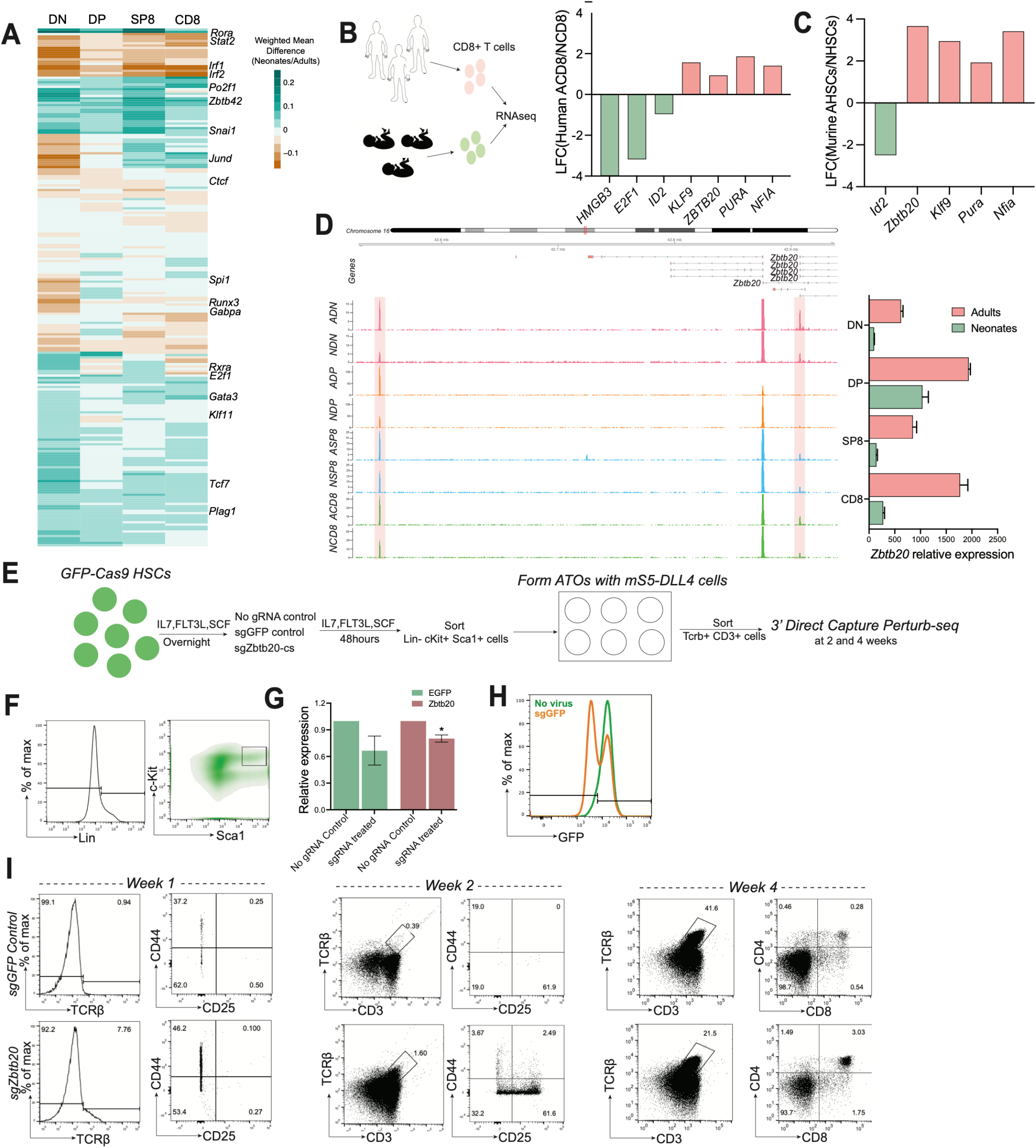
Transcription factors contributing to age-related differences in T-cell development. (A) Significantly differentially active TFs in neonatal versus adult DN, DP, SP8 and CD8+ thymocytes, analyzed by diffTF (Fisher’s exact test, FDR adjusted p<0.05). (B) (Left) Schematic for RNAseq in human adult peripheral CD8+ T cells and human cord-blood CD8+ T cells. (Right) Age-dependent TFs’ expression changes (log2fold change; LFC) in human adult peripheral CD8+ T cells versus human cord-blood CD8+ T cells (Wald test, FDR adjusted p <0.05). (C) Age-dependent TFs’ expression changes (log2fold change; LFC) in murine adult bone marrow HSCs (AHSCs) relative to fetal liver HSCs (NHSCs, FDR adjusted p <0.05). (D) (Left) ATAC-seq signal tracks of the indicated gene in adult and neonatal DN, DP, SP8 and CD8+ T-cell stages. Tracks are group-scaled per developmental stage. (Right) Normalized expression of *Zbtb20* in adult and neonatal thymocytes from indicated stages. (E) Experimental design to test how the loss of *Zbtb20* in ATOs affects T cell development. (F) Representative flow plots display expression levels of HSC markers; lineage, Sca1 and c-Kit levels. (G) Relative expression of EGFP and Zbtb20 in control HSCs and HSCs that received gRNA against EGFP or Zbtb20. (H) Percentage of EGFP positive population in HSCs that either received or did not receive gRNA against EGFP. (I) Representative flow plots of expression levels of CD3, TCRβ, CD44, CD25, CD4 and CD8 at week 1, week 2 and week 4 of thymocytes from control and *Zbtb20* knockdown conditions.

We next asked which novel age-dependent TFs might regulate phenotypic differences in adult and neonatal thymocyte development (Figure 2B). We reasoned that if age-dependent TFs show conserved expression differences in human thymocytes, they might regulate thymocyte developmental differences between adults and neonates. To this end, we compared the expression of these age-dependent TFs in human adult CD8+ T cells and neonatal CD8+ T cells. Human samples were obtained by FACS isolation from human peripheral blood and human cord-blood. Consistent with the mouse thymocyte expression data, human adult CD8+ T cells show higher expression of *ZBTB20*, *PURA* and *KLF9* relative to neonatal counterparts (Figure 6B and S8A). Likewise, *HMGB3* and *E2F1* show higher expression in neonates in both mouse and human CD8+ T cells (Figure 6B). In contrast, *ID2* shows reduced expression in adult CD8+ T cells, which is discordant with our observations in mice (Figure 6B). These findings suggest that age-related expression differences in *ZBTB20*, *PURA*, *KLF9, HMGB3* and *E2F1* are conserved between human and mouse CD8+ T cells.

We next asked which amongst these novel conserved age-dependent TFs are differentially expressed in adult and neonatal HSCs to identify TFs whose differences in expression are linked with their originating progenitor cell identity. We found that *Zbtb20*, *Pura* and *Klf9* are expressed at higher levels in adult compared to neonatal HSCs^121^ (Figure 6C). We further asked which amongst these TFs might: 1) have known roles in the regulation of age-related phenotypes such as altered immunological memory formation and 2) function as a transcriptional repressor, such that its reduced expression in neonates might accelerate T cell development, aiding loss of function studies in adult thymocytes. Amongst these TFs*, Zbtb20,* a transcriptional repressor, regulates immunological memory formation in mature CD8+ T cells^84^. Notably, *Zbtb20* has reduced expression and accessibility in neonates across thymocyte development (Figures 6D and S8B). We hypothesized that the reduced expression of *Zbtb20* may play a role in regulating neonatal thymocyte development and its kinetics.

To investigate whether *Zbtb20* regulates thymocyte developmental rates, we used CRISPR to perturb *Zbtb20* in a three-dimensional artificial thymic organoid (ATO) system, which closely recapitulates T cell developmental stages from HSCs through DN, ISP (Intermediate Single Positive), DP to SP8 stages (Figure 6E)^49^. Our strategy was based on the 3’ Direct Capture Perturb-seq approach, which involves CRISPR-based perturbations of *Zbtb20* coupled with single-cell RNA-sequencing readouts (Figure 6E) ^48,122^, a strategy that combines determination of which cells have undergone deletion of the *Zbtb20* together with a readout for the transcriptional response to *Zbtb20* loss.

To compare the T cell developmental rates in control and *Zbtb20* knockdown ATOs, we generated ATOs from EGFP-Cas9 HSCs (Lineage^−^, Sca1^hi^, c-Kit^hi^ HSCs) that either received no gRNA, gRNAs against EGFP (sg*GFP*) or gRNAs against *Zbtb20* (sg*Zbtb20)*, and harvested thymocytes (TCRβ^hi^ and/or CD3^hi^) at three timepoints (Weeks 1, 2 and 4; Figures 6E and 6F). We observed that sg*GFP*-treated and sg*Zbtb20*-treated EGFP-Cas9 HSCs had reduced expression of EGFP and *Zbtb20*, respectively (Figures 6G and 6H). By Week 1, loss of *Zbtb20* modestly accelerated T cell development (Figure 6I). Strikingly, by Week 2, loss of *Zbtb20* resulted in 32% of thymocytes reaching DN4 compared to only 19% in controls (Figures 6I and S8C). This difference in developmental progression was also observed at Week 4 (Figures 6I and S8C). These observations suggest that loss of *Zbtb20* in adult thymocytes accelerates thymocyte development, explaining, in part, the accelerated thymocyte development characteristic of neonatal thymocytes, which express *Zbtb20* at low levels (Figure 6D).

### *Zbtb20* regulates gene regulatory programs that affect T-cell development

We next investigated how the loss of *Zbtb20* might impact gene regulatory programs that control T-cell development. We hypothesized that *Zbtb20* loss would increase the expression of transcriptional targets that promote thymocyte differentiation. To identify transcriptional targets affected by *Zbtb20* loss, we compared the transcriptional responses between thymocytes that lost *Zbtb20* (sg*Zbtb20*) and control thymocytes (no gRNA and sg*GFP*). For this purpose, we performed 3’ Direct Capture Perturb-seq on thymocytes (CD3^hi^ and TCRβ^hi^) from ATOs cultured to two different time points, Week 2 (T1) and Week 4(T2)^48,122^.

The first two dimensions in uniform manifold approximation and projection (UMAP) of the transcriptomes reflected separation in developmental states, with 7 clusters identified (Figure 7A). We utilized differentially expressed genes in any given cluster relative to all the clusters to identify representative markers (Figures 7B, 7C and S9A). In both control and sg*Zbtb20* induced conditions, cells from clusters 0, 1, 4, 5 and 6 expressed DN1 to DN3 transition genes, with many of these clusters appearing at Week 2 (Figures 7B, 7C and S9A). Genes associated with T cell lineage commitment, intermediate SP stage (ISP), and DP stage were expressed in clusters 3 and 2, respectively, which appear predominantly at Week 4 (Figures 7B and 7C). We further validated the developmental progression through pseudo-time analysis, which predicts the relative temporal progression of cells along a developmental trajectory. This pseudo-time trajectory aligned closely with actual progression in both control and sg*Zbtb20* conditions at time points T1 and T2 (Figures 7B, 7C and 7D). Correlating the pseudo-time trajectory with the UMAP coordinates reveals that both for controls and *sgZbtb20* conditions, UMAP1 and UMAP2 positions could relate cell states to the normal developmental progression.

**Figure 7.**
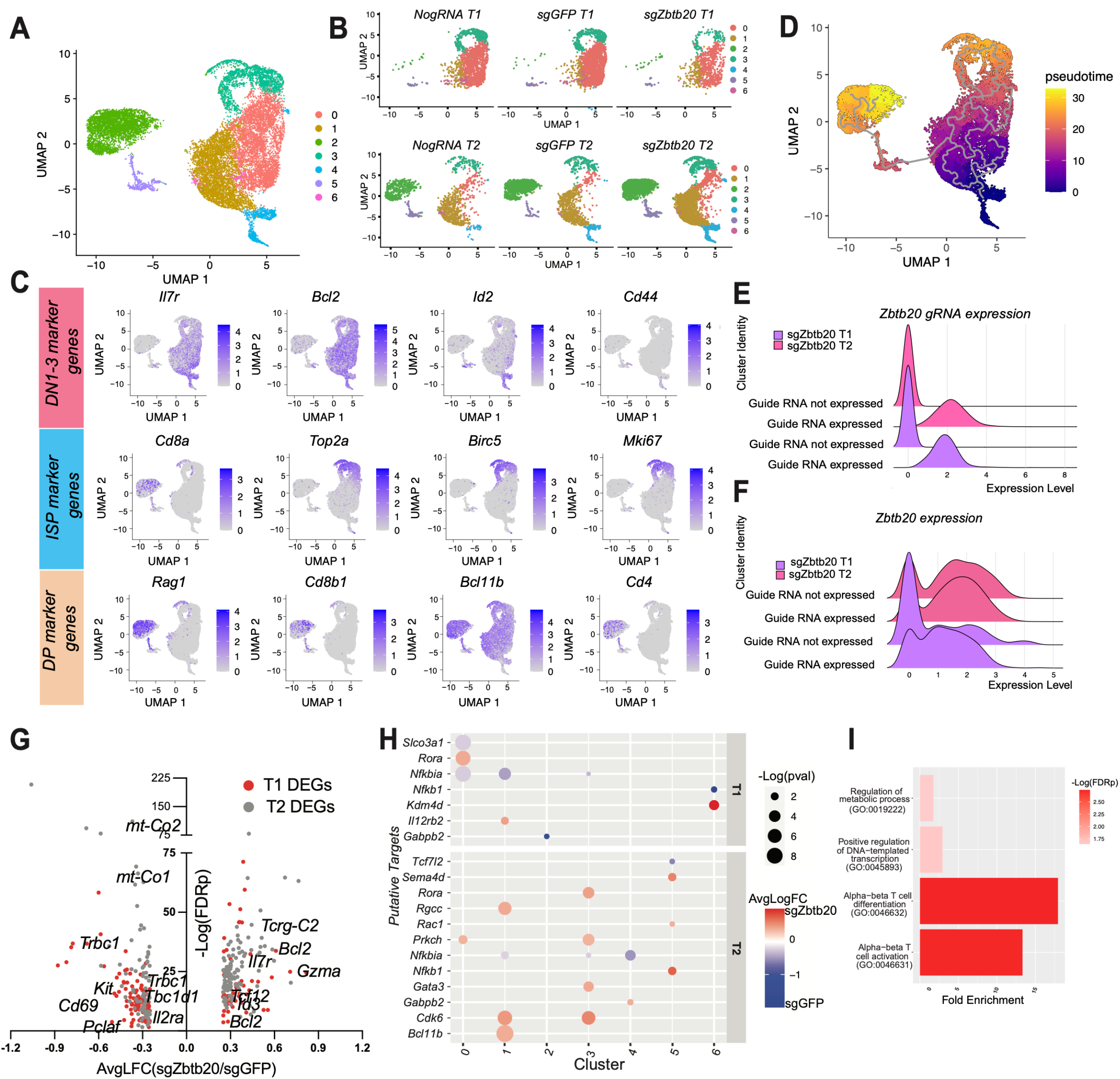
Zbtb20 changes single cell-transcriptomics. (A) UMAP of scRNA-seq profiles of control and *Zbtb20* knock-down ATOs, aggregated across all samples. (B) UMAP of scRNA-seq profiles of ATOs that received no gRNA and gRNAs against EGFP or *Zbtb20*, each at two time points (2 and 4 weeks, T1 and T2, top and bottom, respectively). (C) Color intensity in UMAP displays expression levels of indicated genes, which represent different T cell development stages. (D) The pseudotime score of each cell is displayed in UMAP by color. (E) Ridgeplots comparing expression levels of *Zbtb20* gRNA in clusters that have or do not have *Zbtb20* gRNA expression. (F) Ridgeplots comparing expression levels of *Zbtb20* transcript in clusters that have or do not have *Zbtb20* gRNA expression. (G) Volcano plots compare average log2 fold-changes (LFC) of significant DEGs across all clusters in cells that have *Zbtb20* gRNA expression relative to the control condition at two time points (p<0.05). (H) Dot plots compare average log2 fold-changes of significantly changing putative targets at indicated clusters in cells that have *Zbtb20* gRNA expression relative to the control condition (p<0.05). (I) Significant GO biological processes in significantly changing putative targets, analyzed by PANTHER overrepresentation test (Fisher’s exact test, FDR adjusted p<0.05), focusing on relevant and repetitive GO biological processes.

To compare the transcriptome profiles of *Zbtb20* knockdown versus control conditions, we identified the subset of cells that received *Zbtb20* gRNA (Figures 7E and S9B). We observed a modest decrease in the expression of *Zbtb20* in cells that received *Zbtb20* gRNA (Figures 7E and 7F). Analyzing these *Zbtb20* gRNA-expressing cells, we identified differentially regulated genes between them and control conditions across all clusters at time points T1 and T2. Notably, T cell developmental landmark genes, including *Trbc1*, *Kit, Il2ra, Tcrg-C2, Tcf12* and *Bcl2* were altered upon *Zbtb20* loss relative to control conditions. Consistent with its previously known roles, T cell activation (*Cd69* and *Il7r)* and mitochondrial metabolism regulation (*mt-Co2* and *mt-Co1)* genes showed expression changes upon *Zbtb20* loss (Figure 7G)^84,123^. Thus, many T cell developmental genes, activation genes, and mitochondrial metabolism genes responded to perturbations of *Zbtb20* levels (Figure 7G).

Next, to define direct *Zbtb20* putative targets that were affected by the loss of *Zbtb20* and that may influence T cell developmental rates, we enriched for differentially expressed genes that harbor a *Zbtb20* binding motif within their associated accessible chromatin regions (Figures 7H, 7I, and S9C). Interestingly, genes involved in T cell commitment, including *Bcl11b* and *Gata3,* show increased expression upon *Zbtb20* loss (Figures 7H,7I, and S9C). Their increased expression implies that *Zbtb20* acts as a transcriptional repressor, regulating the expression of *Bcl11b and Gata3*. Furthermore, consistent with previous known roles of *Zbtb20* in regulating *NF-κB* signaling, *NF-κB* signaling components, including *Nfkbia*, *Nfkb1* and *Rora,* were altered upon *Zbtb20* loss (Figures 7H, 7I, and S9D)^124^. *NF-κB* signaling has well-known roles in the regulation of cell survival and fitness. These results suggest how *Zbtb20* may regulate cell fitness and accelerate T-cell developmental rates (Figures 7H, 7I and S9D).

Overall, loss of *Zbtb20* affects T cell developmental genes, *NF-κB* signaling components, T cell activation and metabolism genes. Reduced endogenous *Zbtb20* expression in neonates during early thymocyte development, along with other age-dependent TFs, might accelerate its T cell development rates. These results suggest that *Zbtb20* is a transcriptional regulator of age-related differences during thymocyte development.

## DISCUSSION

Adult and neonatal thymocytes originate from different anatomical niches and exhibit markedly distinct differentiation kinetics, resulting in mature T cells with profoundly different properties, including distinct activation dynamics, memory potential and responsiveness to innate stimulation^1,69^. Many of these differences are associated with gene regulatory changes within mature naïve CD8+ T cells. Here, we set out to determine when these regulatory changes are established. We observed both early divergence in T cell differentiation programs, consistent with their distinct developmental origin (Figure 2B), as well as more gradual divergence in specific gene regulatory programs, possibly due to thymic microenvironmental and anatomical variations (Figures 3A and 3B). We find that the chromatin landscapes are distinct from the earliest stages, presumably shaped by their distinct origin, and are associated with the divergence of these gene expression programs (Figures 4B and 4C). We also observe a transcriptional shutdown during the DP stage, which may impact gene regulatory program differences during thymocyte development (Figure 4E). Strikingly, we observe gene poising as a source of divergence within the age-related gene expression programs (Figure 5E). These findings highlight the ontogeny of divergent gene regulatory programs between adults and neonates. Below, we discuss how our findings shed light on age-related differences in phenotype and function.

We identify a gene module that highlights differences in gene expression programs between adults and neonates from the earliest stages of genesis. In the age-dependent gene module, cell cycle regulation and chromatin organization genes are upregulated in neonates while effector response genes and TCR recombination genes are upregulated in adults (Figure 2B). Neonatal thymocytes preferentially express multiple cell cycle genes at the DN and the SP8 stages, aligning with their increased proliferative ability during those developmental stages (Figures 2B-2E). The increased neonatal cell cycle transcriptional regulation may be linked to the higher proportion of cycling cells among HSCs from the fetal liver, which become neonatal thymocytes^71,121^. However, cell cycle gene divergence is less pronounced during the DP stage, and the mechanism behind this reduced divergence is unclear. Furthermore, as cell cycle programs are coupled with differentiation programs, the increased neonatal cell cycle transcriptional regulation may also regulate the accelerated differentiation kinetics observed in neonatal thymocyte development^125,126^. In contrast to the cell cycle programs upregulated in neonates, effector response genes such as *Id2*, *Bach2* and *Il7r* are upregulated in adults across thymocyte development (Figures 2B, 2C, 2F-2I and 3A). *Il7r* contributes to the long-term survival of CD8+ T cells. As such, reduced *Il7r* in neonates may inhibit their potential to form long-lived memory cells^99^. Similarly, since *Bach2* regulates the differentiation of memory CD8+ T cells, reduced *Bach2* expression in neonates may inhibit their potential to differentiate into memory CD8+ T cells^74–76,82^. Likewise, *Id2* expression is higher in adult thymocytes during thymocyte development, while *Id3* is higher in neonates (2F-2I). Despite *Id2* expression being discordant between human and mouse adult and neonatal thymocytes, these expression differences suggest a possible role in the regulation of both effector response and cell cycle programs between adults and neonates (Figure 6B). Additionally, the age-dependent gene module includes *Dntt*, which is expressed at lower levels in neonates, consistent with previous findings (Figure 2B)^3,127^. The resulting reduced neonatal TCR diversity might impact pathogen recognition in neonates. Overall, we propose this age-dependent gene module as a resource for dissecting programs that change between adults and neonates across thymocyte development.

We also investigated how chromatin landscapes differ between adult and neonatal thymocytes. Neonates exhibit greater chromatin accessibility during early thymocyte development, while adults show increased accessibility in later stages (Figure 4B). These changes correlate with shifts in gene expression, particularly in effector response genes such as *Bach2*, *Il7r* and *Cd69*, which are upregulated in adults, and *E2f1*, *Ccnb1* and *Cenph*, cell cycle genes, which are upregulated in neonates (Figures 4B-4D). However, several age-related gene expression programs lack corresponding changes in accessibility, suggesting that alternative gene regulatory mechanisms contribute to these programs. Notably, neonates demonstrate a higher prevalence of poised genes compared to adults, governing distinct gene expression programs (Figures 5A-5D). Adults display poised *NF-κB* signaling components, while neonates harbor poised innate immune response genes (Figure 5E). This poising may contribute to the differences in innate immune responsiveness between adult and neonatal CD8+ T cells. Together, we put forth the model that distinct chromatin states in neonates and adults, potentially influenced by their disparate origins, regulate the transcriptional disparities observed between them.

Given the age-related differences in chromatin accessibility, we identify TFs that are involved in generating age-related differences in gene expression programs (Figure 6A). Notably, TFs such as *Gata3*, *Tcf7, Ctcf* and others show differential activity during adult and neonatal thymocyte development (Figure 6A). *Zbtb20*, a conserved transcriptional repressor, displays higher chromatin accessibility and expression in adult HSCs and thymocytes relative to neonatal HSCs and thymocytes (Figures 6B and 6C). We find that loss of *Zbtb20* leads to accelerated T cell developmental kinetics during adult thymocyte development (Figure 6I). Furthermore, the loss of *Zbtb20* derepresses key T cell developmental genes, including *Bcl11b* and *Gata3* (Figure 7H), both of which harbor *Zbtb20’s* binding motif. Our data suggest that *Zbtb20* binds to regulatory regions of *Bcl11b* and *Gata3* and may recruit corepressor proteins to repress target genes^124,128^. Finally, consistent with previous results, our data is indicative of *Zbtb20* altering *NF-κB* mediated signaling (Figure S9D), and this may improve cell survival and impinge on differentiation kinetics. These results underscore *Zbtb20* as a transcription factor that regulates age-related differences in adult and neonatal thymocyte development.

In summary, our study provides a detailed transcriptional and chromatin accessibility map for adult and neonatal thymocyte development. By integrating transcriptomic and epigenetic information from adult and neonatal thymocyte subsets, this atlas provides a foundation to dissect the mechanisms underlying differences in adult and neonatal thymocyte development. Moreover, we uncover *Zbtb20* as a novel regulator of age-related differences in thymocyte progression.

## ACKNOWLEDGMENTS

This work was supported by National Institute of Health awards P50HD076210 (to A.W.G., from National Institute of Child Health and Human Development), R01AI110613, U01AI131348 (to B.D.R and A.W.G., from the National Institute of Allergy and Infectious Disease), R01AI105265 (to B.D.R, from the National Institute of Allergy and Infectious Disease), A.W.G and K.M.S. were supported by U24AI152176. K.M.S was supported by K08AI108870. We also thank the Cornell Center for Animal Resource and Education (CARE) for expert mouse breeding assistance. Cell sorting was done at Cornell University’s Flow Cytometry Facility (RRID:SCR_021740) in the Biotechnology Resource Center (BRC). RNA-seq and ATAC-seq profiling projects were coordinated and performed with the assistance of the Transcriptional Regulation and Expression Facility (RRID:SCR_022532); 10x Genomics libraries were prepared, and all Illumina sequencing was conducted at the BRC Genomics Facility (RRID:SCR_021727) at Cornell University. Human cells were collected and processed at the Golisano Children’s Hospital, University of Rochester Medical Center, as part of the Respiratory Pathogens Research Center (NIAID HHSN272201200005C). We thank all A.W.G and B.D.R lab members for helpful discussions on the manuscript.

## AUTHOR CONTRIBUTIONS

D.G.S planned and performed experiments, analyzed and interpreted data, and wrote the manuscript. H.Z analyzed data. V.I.M, I.M.F and N.L performed experiments. Y.J performed experiments and analyzed the data. K.M.S provided intellectual feedback. B.D.R provided intellectual feedback. A.W.G supervised the project and provided intellectual feedback.

## DECLARATIONS

The authors declare no competing issues.

## DATA AND MATERIALS AVAILABILITY

The accession numbers for the RNA-seq, ATAC-seq and scRNA-seq datasets reported here are GSE267922, GSE267918 and GSE267923 respectively. All other data needed to support the conclusions of the paper are present in the Supplementary Materials and with Mendeley data volume, V1, doi: 10.17632/kdxhf7pmmk.1.

## MATERIALS AND METHODS

### Mice

C57BL/6 mice were purchased from Charles River Laboratories and maintained in our colony. At the time of experimentation, adult mice were 8-12 weeks old and neonatal mice were 5-7 days old. All mice were sex-matched within experiments; female mice and male mice were used for RNA-seq, ATAC-seq and staining experiments. EGFP-Cas9 mice were purchased from the Jackson Laboratories (strain #026179), crossed to gBT-I TCR transgenic mice (TCRαβ specific for the HSV-1 glycoprotein B_498-505_ peptide SSIEFARL), and then maintained in our colony. gBT-I transgenic mice were generously provided by Dr. Janko Nikolich-Zugich (University of Arizona, AZ)^129^. Mice were kept under specific pathogen-free conditions at Cornell University College of Veterinary Medicine, accredited by the American Association of Accreditation of Laboratory Animal Care. All experiments were conducted with approval from the Institutional Animal Care and Use Committee at Cornell University.

### Thymocyte and lymphocyte isolation

Thymi and spleens from adult or neonatal mice were collected in RP-10 medium (RPMI 1640 with 10% fetal bovine serum) and mashed through a 40μM cell filter to create a single-cell suspension. Spleens and thymi underwent magnetic enrichment with CD8a and CD4 MicroBeads (Miltenyi Biotec, #130-117-044; #130-117-043, respectively), per the manufacturer’s instructions. The bound thymic fraction was sorted for DP thymocytes, while the eluted fraction underwent CD8a magnetic enrichment as described previously, after which the bound fraction was FACS-sorted (Fluorescence-Activated Cell Sorting) for SP8 thymocytes, while the eluted fraction was FACS-sorted for DN thymocytes. Thymocyte counts were taken by diluting a single-cell thymus suspension and counting on a MoxiZ cell counter.

### Flow cytometry

For thymocyte development samples, single-cell suspensions of thymi/spleens or fractionated thymi were stained in a 500μL antibody cocktail (PBS + 0.5% BSA + 2mM EDTA; all antibodies were used at 1:200 dilution) for 30 minutes at 4°C in the dark. Populations of interest were FACS-sorted (Sony MA900). Id2, Id3 and Ki67 were stained using the Foxp3 staining buffer set (eBioscience, #00-5523-00) according to the manufacturer’s instructions. FACS was conducted using the Symphony A3 (BD Biosciences) and analyzed on FlowJo. For M-ATO phenotyping, single-cell suspensions of M-ATO thymocytes were generated as described below and were stained in 500μL antibody cocktail for 30 minutes at 4°C in the dark. FACS was conducted using the Attune (ThermoFisher Scientific). The following antibodies were used: CD8a (53-6.7), CD4 (GK1.5 or RM-5), gamma delta T cell receptor (eBioGL3), NK1.1 (PK136), CD25 (3C7 or PC61), Ter119 (TER-119), CD44 (IM7), CD45.2 (104). Dead cells were stained with eBioscience Fixable Viability Dye eFluor 780 (ThermoFisher Scientific, #65-0865-14) at 1:1,000 dilution.

### Human samples

Frozen de-identified full-term cord blood (39 to 41 weeks; neonatal) or adult (23 to 32 years) peripheral blood mononuclear cells (CBMCs or PBMCs, respectively) were obtained from the Biorepository Core at the University of Rochester. All study procedures were approved by the University of Rochester Medical Center Internal Review Board (protocol nos. RPRC00045470 and 37933), and all individuals’ caregivers provided informed consent. Individuals were excluded for known congenital anomalies or genetic disorders present or suspected, individual nonviability, and for maternal HIV or inherited disorders known to affect immune development. Frozen CBMCs and PBMCs were quick-thawed and rested overnight in RP-10 medium at 37°C. After resting, adult and neonatal naïve CD8+ T cells were isolated by FACS as CD14^-^CD56^-^CD3^+^CD4^-^ CD8a^+^CD45RO^−^CD45RA^+^CCR7+ from PBMCs and CBMCs respectively.

### Plasmid DNA and cloning

Retroviral vectors containing either sgGFP-constant region (Cr2) or sgZbtb20-constant region (Cr3)-with-capture sequence (cs2) were generated as described below. The constant regions (Cr2 and Cr3) with capture sequence(cs2) were derived from perturb-seq vectors, which allow for 3**’** Direct Capture. Briefly, MSCV-IRES-Thy1.1-DEST vectors^82^ were cleaved using MluI and SalI. Next, the mU6-gRNA-Constant-region (Cr)-with-capture-sequence(cs) module, derived from perturb-seq vectors pMJ179 and pBA900, was integrated into the cleaved vector via Gibson cloning (New England Biolabs, #E2621L) to produce a retroviral vector that contained either mu6-sgGFP-Cr2 (that harbored no capture sequence) or mu6-sgZbtb20-Cr3-cs2 regions ^48,130,131^. Subsequently, these vectors were used for retroviral transductions, as described below.

### HSCs extraction and culture

HSCs were extracted from EGFP-Cas9-gBI+ mice following cervical dislocation. Femur and tibia bones were isolated and flushed with 10ml RP-10 using a 27-gauge needle. Bone marrow extracts were processed into a single-cell suspension and cells from a single mouse were pooled. After pelleting at 500*g* for 5 minutes, cells were resuspended in ice-cold MACS buffer through a 70μM filter. This wash step was repeated twice and cells were pelleted at 500*g* for 5 minutes. Biotin cocktail (1ml) containing CD5 (53-7.3), CD19 (eBio1D3), CD11b (M1/70), GR-1 (RB6-8C5), B220 (RA3-6B2) and Ter119 (TER-119) was added to each pellet and incubated at 4°C for 20 minutes. Following incubation, 9ml MACS was added and spun at 500*g* for 5 minutes. The supernatant was aspirated, and the pellet was resuspended in 750μl streptavidin mix, which contained 75μl streptavidin beads (Miltenyi Biotec, #130-048-101) and 675μl MACS. This mixture was further incubated for 15 minutes at 4°C. After the incubation, 9ml of MACS was added and spun at 500*g* for 5 minutes. After a further spin with 9ml MACS, cells were passed through an LS column (Miltenyi Biotec, #130-042-401) following a buffer wash. Finally, cells were resuspended in transplant media comprising RP-10 and cytokines (mSCF, 50ng/ml, #250-03; IL-6, 10ng/ml, #216-16; IL-3, 10ng/ml, #213-13; and Flt3l, 10ng/ml, #250-31) with 1x penicillin/streptomycin and were cultured overnight in non-tissue culture treated 6 well-plates.

### Retroviral transduction

20 μg of retroviral plasmid DNA (sgGFP-Cr2 or sgZbtb20-Cr3-cs2) with 6 μg pCL-Eco plasmid DNA were transfected using 60 μl Lipofectamine 2000 (Invitrogen, #1668019) in 3 ml OptiMEM (Invitrogen, #31985062) for 8 hours in antibiotic-free media into Platinum-E ecotropic packaging cells (Cell Biolabs, #RV-101). These cells were plated at a concentration of 2 million cells per 10-cm plate two days before transfection. Media in platinum-E ecotropic packaging cells were replaced 7 hours after transfection and these cells were further incubated for 72 hours. Following the incubation, filtered retroviral supernatants were harvested and applied to HSCs (extracted as described above) in plates coated with Retronectin (20 μg/ml; Takara Bio; *#*T100B). The transduction process involved spinning at 2,000*g* for 2 hours at 32°C. Transduced cells were harvested after 48 hours, and Lin^-^Sca^+^Kit^+^ HSCs were isolated after staining and FACS-sorting (Sony MA900), characterized by the following cell-surface antibody phenotyping: Lin^-^ (Ter119^-^, TER119; B220^-^, RA3-6B2; Gr1^-^, RB6-8C5; NK1.1^-^, PK136; CD3^-^, 145-2C11), Sca^+^ (D7) and Kit^+^(2B8).

### Murine artificial thymic organoid (M-ATO) cultures

M-ATOs were generated with modifications as described below^49^. MS5-mDLL4 cells were harvested by trypsinization and resuspended in a serum-free M-ATO RPMI-B27 culture medium. RPMI-B27 media contains RPMI 1640, 4% B27 supplement (ThermoFisher Scientific, #17504-044), 30 μM L-ascorbic acid 2-phosphate sesquimagnesium salt hydrate (Sigma-Aldrich, #A8960-5G; reconstituted in PBS), 1% penicillin/streptomycin, 1% Glutamax, 5 ng/ml rmFLT3L (Peprotech, #250-31L), 5 ng/ml rmIL-7 (Peprotech, #217-17), 10 ng/ml rmSCF (SCF was added only for the first week of culture; Peprotech, #250-03) and β-mercaptoethanol (0.05mM; Sigma-Aldrich, #M7522). RPMI-B27 was made fresh weekly. 1.5x10^5^ MS5-mDLL4 cells were combined with purified transduced Lin^-^Sca^+^Kit^+^ HSCs (1,000 cells per ATO; extracted as described above) and centrifuged at 300*g* for 5 min at 4°C in a swinging bucket centrifuge. Supernatants were carefully removed, and the cell pellet was resuspended in 5 μl RPMI-B27 per M-ATO and mixed by brief vortexing. M-ATOs were plated on a 0.4 μm Millicell transwell insert (EMD Millipore, #PICM0RG50) and placed in a 6-well plate containing 1 mL RPMI-B27 per well. Medium was changed every 3-4 days by aspiration from around the transwell insert, followed by replacement with 1 mL with fresh RPMI-B27/cytokines. M-ATO cells were harvested by adding FACS buffer (PBS/0.5% bovine serum album/2mM EDTA) to each well and briefly disaggregating the M-ATO by pipetting with a 1 mL pipet, followed by passage through a 50 μm nylon strainer. For M-ATO phenotyping, M-ATO thymocytes were isolated after staining (described above) and FACS-sorting at different timepoints (week1, week 2, week 3 to 4). They were then characterized by the cell-surface antibody phenotyping as follows: thymocytes (Live^+^CD45.2^+^CD3^+^TCRβ^+^), DN1-4 (CD3^+^/TCRβ^+^CD44^hi/lo^CD25^hi/lo^) and DP (CD3^+^TCRβ^+^CD4^+^CD8^+^). For single-cell RNA sequencing using 3’ Direct capture perturb-seq approach, M-ATO thymocytes were isolated by FACS-sorting at different time points (Week 2: day 12, Week 3 to 4: day 27) and selected for Live^+^CD45.2^+^CD3^+^TCRβ^+^ populations.

### RNA-seq libraries preparation and sequencing

To extract RNA, sorted thymocyte subsets were resuspended in 1ml of Trizol (ThermoFisher Scientific, #15596018). 200μl of chloroform was added to 1ml of trizol lysates. After vigorous shaking, samples were centrifuged at 16,000*g* at 4°C. During the spin, phase lock gel tubes were prepared by a pre-spin at 12,000*g* at room temperature. 600μl chloroform was added to these pre-spun tubes. The aqueous phase of the trizol-chloroform mixture was added to the phase lock tube containing chloroform. After vigorously shaking, these phase lock tubes were centrifuged at 12,000*g* at 4°C. The aqueous phase from the phase lock tubes was added to isopropanol and glycoblue-containing precipitation tubes. These precipitation tubes were mixed, incubated for 1 hour, and spun at 16,000*g* at 4°C. The supernatant was discarded, and the pellets were washed with ice-cold ethanol. Pellets were further centrifuged at 16000*g* at 4°C. After the aspiration of the supernatant, the pellet was air-dried and resuspended in RNAase-free water. RNA sample quality was confirmed using a Qubit (RNA HS kit; ThermoFisher Scientific) to determine the concentration and the Fragment Analyzer (Agilent) was utilized to determine RNA integrity. PolyA+ RNA was then isolated with the NEBNext Poly(A) mRNA Magnetic Isolation Module (New England Biolabs). TruSeq-barcoded RNAseq libraries were generated with the NEBNext Ultra II Non-Directional RNA Library Prep Kit (New England Biolabs). Each library was quantified with a Qubit (dsDNA HS kit; ThermoFisher Scientific) and the size distribution was determined with a Fragment Analyzer (Agilent) prior to pooling. Libraries were pooled and sequenced on a NovoSeq 6000 instrument in a 2x150 cycle run. On average 21 million mapped reads were generated per library.

### ATAC-seq library preparation and sequencing

The ATAC-seq libraries were prepared based on the Omni-ATAC-seq protocols^132^ with minor modifications. Briefly, 100,000 FACS-sorted cells were used to prepare nuclei. The transposition reaction was performed by incubating 25,000 nuclei with Nextera Tn5 transposase (Illumina) at 37°C for 1 hour. The transposition mixture was purified using a DNA Clean and Concentrator kit (Zymo, #D4004). The ATAC libraries were amplified for 11 cycles using NEBNext 2X MasterMix and Nextera Index primers (New England Biolabs, # M0541S). The amplified libraries were size selected using AMPure beads (Beckman Coulter, #A63880) to remove large DNA fragments. Libraries were pooled and sequenced on a NovoSeq 6000 instrument in a 2x150 cycle run. On average 29 million mapped read pairs were obtained per library.

### 3’ Direct-capture perturb-seq library preparation and sequencing

Single-cell ATO thymocyte suspensions were run on a Chromium X instrument and libraries were prepared following the user guide: Chromium Next GEM Single Cell 3’ RNA-Seq Assay v3.1 Dual Index with Feature Barcode technology for CRISPR Screening (10x Genomics, #CG000316, RevD)^133^. We targeted 6,000 cells and used 12 cycles of cDNA amplification. Sample quality was confirmed using a Qubit (RNA HS kit; ThermoFisher Scientific) to determine concentration and a Fragment Analyzer (Agilent) to determine RNA integrity. Every sample had an associated feature barcode except the control libraries. Libraries were sequenced on a NovoSeq X instrument in a 2x150 cycle run. On average 64,000 reads were generated per cell.

### RNA-seq data processing and analysis

The paired-end reads (from four independent biological replicates for each developmental stage and age) were processed using trim galore and fastq files were aligned using STAR 2.7.10a with mm10 mouse genome assembly ^134^. To assess gene body coverage, a gene body coverage python package (RseQC geneBody_coverage.py) was utilized. Reads mapped to annotated features were counted using featureCounts^135^. The Pearson correlation coefficient (r) was used to compare the correlation between biological replicates. DEseq2 was used to normalize the raw counts and identify differentially expressed genes^136,137^. The principal components analysis (PCA) plot was generated using plotPCA function (top 500 genes) within DESeq2. Differentially expressed genes (DEGs) from RNAseq data were identified using adjusted p-value < 0.05 and log_2_-fold change (LFC) > 0.589. All plots with RNA expression are shown as normalized and log_2_ transformed (unless stated).

### Identification of age-dependent gene module

To identify the age-dependent gene module, adults and neonates were compared within every developmental stage. Significantly changing DE genes (adjusted p-value < 0.05 and LFC > 0.589) at an individual stage were utilized and intersected to identify the 275 genes in the age-dependent gene module.

### Identification of developmental stage-specific signature genes

Adults and neonates were compared across multiple developmental stages; age and developmental stage were included in the model formula to adjust for age-and stage-related statistical variations. To define time point specific signature genes, we performed hierarchical clustering on DEGs and split the dendrogram to obtain clusters that show high expression at every developmental timepoint.

### Functional annotation analysis

Functional annotation analysis was done either using R enrichGO () from clusterProfiler or GO-SLIM PANTHER overrepresentation test (Fisher’s Exact test, all the displayed terms have FDR adjusted p<0.05 unless stated)^137^. Statistically significant GO (biological processes) were identified; only repeated and relevant GO biological processes are shown and discussed in the figures. Repetitive and relevant GO terms were curated manually and validated with a language model^138,139^ (the complete list of GO terms is included as supplementary tables; Tables S4 and S6). For functional annotation using ATAC-seq ACRs, GREAT (Genomic Regions Enrichment of Annotations Tool) analysis was used (default parameters to calculate the genes associated to ATAC–seq peaks and associated GO terms were chosen)^140^. GO terms that suggest temporal correlation between stage-specific genes and peaks are discussed.

### ATACseq data processing and analysis

Fastq files were aligned to the mouse mm10 reference genome with bwa. Unmapped, unpaired, and mitochondrial reads were removed using samtools. PCR duplicates were removed using Picard. Peak calling was performed on four replicates with MACS2 with an FDR q-value < 0.05. A union peak list of each data set was created by combining all peaks in all samples, merging overlapping peaks using MACS2 and keeping peaks that were called in more than one sample as a .saf file. The number of reads in each peak was determined with featureCounts subread package^135^. Peaks were annotated using HOMER. Each peak was assigned to the nearest gene based on the shortest distance between the peak and the gene’s promoter on either strand; promoters were defined as 1 kb upstream and 100 bp downstream of annotated transcription start site (TSS). Each gene was assigned to a peak cluster with the greatest number of gene associated peaks; for ties, genes were randomly assigned to one of the tied clusters. The principal components analysis, Pearson correlation coefficient computation, and comparison of adult and neonatal stage-specific peaks were computed as mentioned in the RNAseq data processing section. Differentially accessible chromatin regions (DACRs) were identified using adjusted p-value < 0.05 and LFC > 0.1. All the ATAC-seq peaks were normalized and log_2_ transformed.

### Clustering

All clustering was performed with pheatmap from the pheatmap R package or ComplexHeatmap R package (adjusted p-value ≤ 0.05, LFC > 0.589 for RNAseq and LFC > 0.1 for ATACseq). When the ComplexHeatmap package was used, counts were converted to z-scores across samples (row-wise z-scores) prior to clustering analysis. In the pheatheatmap, counts were scaled by z-scores within the function (default methods were used). The k for row/column cluster identification was chosen based on the biological question as indicated.

### Identification of poised genes

To identify early differentially accessible peaks in adults and neonates, DN ACRs were compared to DP ACRs (DN to DP, adjusted p < 0.05, positive LFC > 0.1). Early, mid and late differentially expressed genes in adults and neonates were defined by comparing DP genes to DN genes (DP/DN), SP8 genes to DP genes (SP8/DP), and CD8 genes to SP8 genes (CD8/SP8) respectively (adjusted p < 0.05, positive LFC >0.589). Poised genes were identified by subtracting early DEGs from early DACRs in adults and neonates, then intersecting this subset with mid and late DEGs from adults and neonates. Two subsets were obtained, indicating poised genes for mid-and late-development in adults and neonates, respectively. Common poised genes were subtracted from this subset to identify unique poised genes specific to adults and neonates. Gene Ontology (GO) analysis was performed as described earlier and terms that explain temporal regulation of ACD8 or NCD8 signature genes are shown in the figures.

### *Zbtb20* putative targets/TF motif enrichment

*Zbtb20* putative targets were identified and analyzed using the HOMER tool *findMotifsGenome.pl* [findMotifsGenome.pl -size given -find Zbtb20Motif]. JASPAR^141^ maintains a custom database of known motifs based on experimentally defined data sets. From the set of known motifs, we filtered for *Zbtb20* motifs. Because many members of a TF family share a similar core motif, we chose one representative *Zbtb20* family motif (UN0143.1). We further converted this JASPAR motif to a HOMER-compatible format for analysis using HOMER.

### Differential TF activity

Quantification of differential TF activity was performed by DiffTF^117^. Comparison was performed between adult and neonatal samples at each stage (DN, DP, SP8, CD8). Analytical mode was used as recommended for comparisons between conditions with 3 replicates (parameters: nPermutations = 0, nBootstraps = 1000). Bam and narrowPeak files from ATAC-seq and count tables from RNA-seq were used as input for DiffTF. TF Motifs from HOmo sapiens COmprehensive MOdel COllection (HOCOMOCO) v10 were used, with the addition of motifs of four age-dependent TFs of which the human motifs were found in JASPAR^142^ (*Zbtb20*: UN0143.1, *Zbtb42*: UN0312.1, *Klf9*: MA1107.1, *Klf11*: MA1512.1). For the four TFs, the position weight matrices of their human motifs were downloaded from JASPAR, and fimo^141^ was used to identify their occurrences in the mouse genome with default parameters.

### Single-cell RNA-seq data processing and analysis

Fastq files were generated with cellranger mkfastq (10x Genomics) by NovoSeq X. Raw count tables were generated with cellranger count v7.2.0 (10x Genomics) [cellranger count --id = ID --transcriptome = /path/to/ refdata-gex-mm10-2020-A/--fastqs = /path/to/directories --sample = list --feature-ref= /path/to/feature-ref-file/ -- localcores=16 -check-library-compatibility= false]. The feature reference files used an untethered approach [pattern =(BC) sequence = GCTCACCTATTAG/capture sequence]. To create aggregate files, cell ranger aggr was utilized [cell ranger aggr --id = ID –csv=/path/to/molecule.h5 files/]. The aggregated file of control and *Zbtb20* loss conditions were utilized for downstream analysis. Data normalization, cell clustering, and differential expression were carried out using the Seurat R package v4.3.0^143^. Cells with more than 10% of mitochondria genes were excluded from the analysis. Read counts were normalized using the Seurat NormalizeData function. Variable features for downstream analysis were identified using the Seurat FindVariableFeatures function (nfeatures = 2000). Data was then scaled and centered using ScaleData function in Seurat. PCA was performed using the Seurat RunPCA function. Uniform Manifold Approximation and Projection (UMAP) and Graph-based clustering (using the Seurat FindNeighbors and FindClusters functions) were performed with the top 10 principal components at the resolution of 0.15. Trajectory analysis was performed with Monocle 3^144^, with cells from all sample types and timepoints as input. The trajectory was learned by order_cells function and plotted by plot_cells function in R package monocle3. Differential expression analysis was performed comparing cells of the sgGFP condition and cells of the sgZbtb20 condition with gRNA targeting *Zbtb20* expressed. The analysis was conducted for samples at time points 1 and 2 separately using the FindMarkers function in Seurat with default parameters.

## SUPPLEMENTAL INFORMATION

**Figure S1 Isolation of Adult and neonatal thymocytes and splenocytes, related to Figure 1**.

(A) Flow cytometry analysis of indicated markers in pre-and post-sorted samples for indicated cell types.

(B) Absolute cell numbers of adult (red bars) and neonatal (green bars) DN1, DN2 and DN3 thymocytes, compared with two-way ANOVA (ns, not significant; *, *p* < 0.05; ** *p* < 0.01).

(C) Absolute cell numbers of adult and neonatal DP thymocytes; otherwise, as described in panel B.

(D) Absolute cell numbers of SP8 thymocytes post-sort; otherwise, as described in panel B.

**Figure S2 Isolation of Adult and neonatal thymocytes and splenocytes, related to Figure 1**.

(A) Raw and normalized counts distribution in various adult and neonatal transcriptomes obtained using RNA-seq.

(B) Raw and normalized raw counts distribution in various adult and neonatal thymocyte accessible chromatin regions obtained using ATAC-seq (ADN4, ADP4, ASP84, ACD84, NDN4, NDP4, NSP84 and NCD84 outliers have reduced read counts).

(C) Gene body (5’ to 3’) bias showing ADN1 as an outlier.

(D) Principal component analysis (PCA) of RNA-seq profiles (top 500 genes), with percentage variance associated with each axis indicated. Adult and neonatal samples are denoted by circular and triangular points, respectively, with developmental stage color-coded, as indicated.

(E) Heatmap representing Pearson correlation (*r*) coefficient between adult and neonatal thymocytes.

**Figure S3 Gene regulatory programs that differ between adults and neonates across thymocyte development, related to Figure 2**.

(A) Significant DEGs at a specific thymocyte developmental stage (row normalized expression; z-scores; see color-coded scale bar in adults and neonates, left and right halves, respectively; see Table S1 for full list of genes in age-dependent gene module; Wald test, FDR adjusted p <0.05).

(B) Repetitive, relevant, and significant GO processes, identified by a language model on the full list of significant GO processes in the age-dependent gene module (see Table S2 for the full list of significant processes; FDR adjusted p <0.05).

(C) Cell cycle genes in the age-dependent gene module, highlighting gene names.

(D) *Id2* normalized transcript expression (z-scores) in adult and neonatal thymocytes.

(E) *Id3* normalized transcript expression (z-scores) in adult and neonatal thymocytes.

**Figure S4 Adult and neonatal developmental stage-specific gene expression programs, related to Figure 3**.

(A) Significant DEGs between adults and neonates during thymocyte development, accounting for developmental stage variations (row normalized expression; z-scores; see color-coded scale bar in adults and neonates, left and right halves, respectively; see Tables S3 and S5 for full list of adult and neonatal signature genes; Wald test, FDR adjusted p <0.05).

**Figure S5 Gene regulatory programs that differ between adults and neonates across thymocyte development, related to Figure 3**.

(A) Top to bottom, repetitive, relevant, and significant GO processes, analyzed by a language model on the full list of significant GO processes in ADN, ASP8 and ACD8 signature genes (see Table S4 for the full list of significant GO processes; FDR adjusted p <0.05).

(B) Heatmap of *Spi1* cluster, comparing expression (normalized expression; see color-coded scale bar) in adults (left) and neonates (right).

(C) Top to bottom, Repetitive, relevant, and significant GO processes, analyzed by a language model on the full list of significant GO processes in NDN and NDP signature genes (see Table S6 for the full list of significant GO processes; FDR adjusted p <0.05).

**Figure S6 Chromatin accessibility during thymocyte development, related to Figure 4**.

(A) Heatmap representing Pearson correlation coefficient(*r*) between accessible chromatin regions obtained using ATAC-seq.

(B) Cluster dendrogram representing Euclidean distance between various adult and neonatal accessible chromatin.

(C) Top to bottom, PCA of ATAC-seq profiles (top 500 genes) of adult (red circles) and neonatal (blue triangles) at DN, DP SP8, and CD8+ T stages.

(D) PCA of ATAC-seq profiles (top 500 genes), with percentage variance associated with each axis indicated. Adult and neonatal samples are denoted by circular and triangular points, respectively, with developmental stage color-coded, as indicated.

(E) Accessibility around cell cycle genes in adult and neonatal DN stage (row normalized accessibility; see color-coded scale bar).

**Figure S7 Transcription factors implicated in thymocyte development, related to Figure 6**.

(A) TFs that show increased binding in adult or neonatal DN thymocytes, analyzed by diffTF (Fisher’s exact test, FDR p adjusted < 0.05).

(B) TFs that show increased binding in adult or neonatal DP thymocytes, analyzed by diffTF (Fisher’s exact test, FDR p adjusted < 0.05).

(C) TFs that show increased binding in adult or neonatal SP8 thymocytes, analyzed by diffTF (Fisher’s exact test, FDR p adjusted < 0.05).

(D) TFs that show increased binding in adult or neonatal CD8+ T cells, analyzed by diffTF (Fisher’s exact test, FDR p adjusted < 0.05).

**Figure S8 Transcription factors controlling age-related differences in T cell developmental rates, related to Figure 6**.

(A) Normalized counts of *Zbtb20* in human adult peripheral blood-derived CD8+ T cells and cord blood-derived CD8+ T cells (Wald test, FDR p adjusted < 0.05).

(B) ATAC-seq signal tracks of the indicated gene in adult and neonatal DN, DP, SP8 and CD8+ T-cell stages. Tracks are group-scaled per dataset.

(C) (Left) Percentage of DN thymocyte (CD3^hi^TCRβ^hi^) subsets DN1, DN2, DN3 and DN4 at week 1. (Right) Percentage of DN thymocyte (CD3^hi^TCRβ^hi^) subsets DN1, DN2, DN3 and DN4 at week 2. (Student’s t-test, p < 0.05).

**Figure S9 Impact of Zbtb20 perturbation on gene expression, related to Figure 7**.

(A) Color intensity in UMAP displays expression levels of indicated genes.

(B) Color intensity in UMAP displays expression levels of *Zbtb20* gRNA.

(C) Average expression of the indicated genes in each cluster and the percentage of cells that express the indicated genes in each cluster.

(D) Normalized enrichment scores of the indicated pathway, generated using GSEA, based on *Zbtb20* putative direct targets expression changes relative to controls.

**Table S1: List of genes in the age-dependent gene module, related to** Figure 2.

**Table S2: List of significant GO biological processes in the age-dependent gene module, related to** Figure 2.

**Table S3: List of ADN, ADP, ASP8 and ACD8 signature genes, related to** Figure 3.

**Table S4: List of significant GO biological processes in ADN, ASP8 and ACD8 signature genes, related to** Figure 3.

**Table S5: List of NDN, NDP, NSP8 and NCD8 signature genes, related to** Figure 3.

**Table S6: List of significant GO biological processes in NDN and NDP signature genes, related to** Figure 3.

