## Supplementary figures and images for "Gene Regulatory Programs that Specify Age-Related Differences during Thymocyte Development"

### Supplemental_Figures

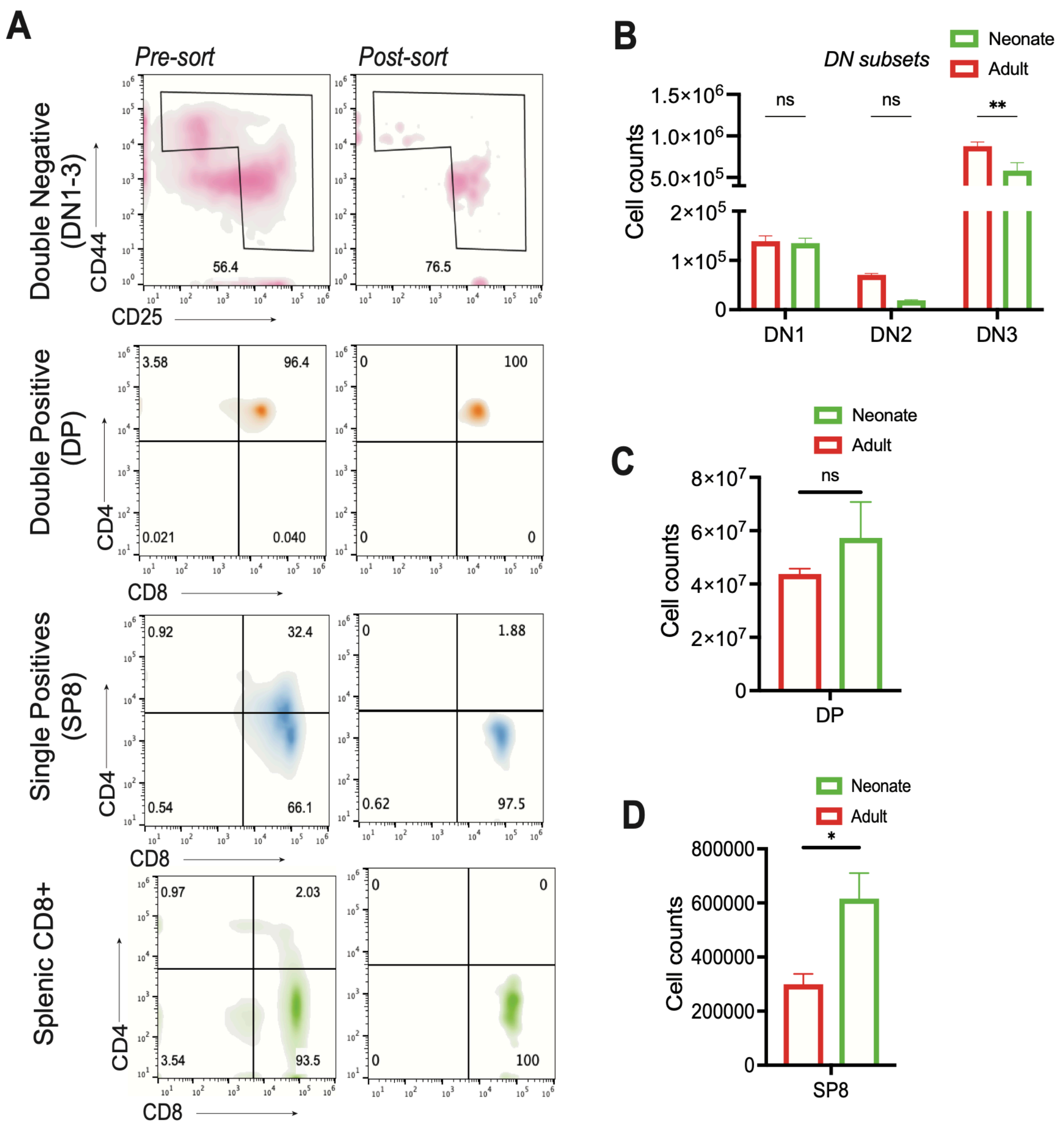

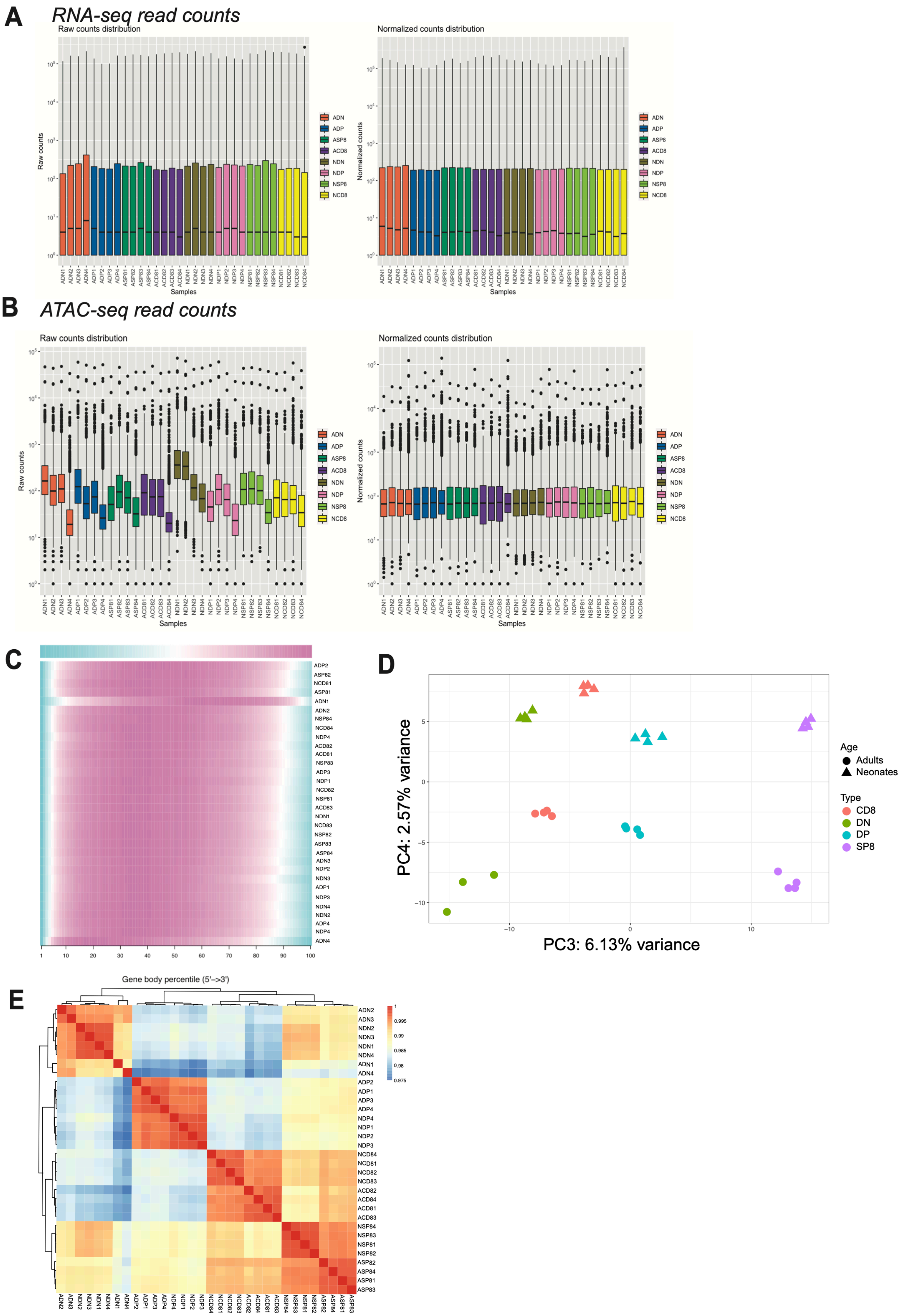

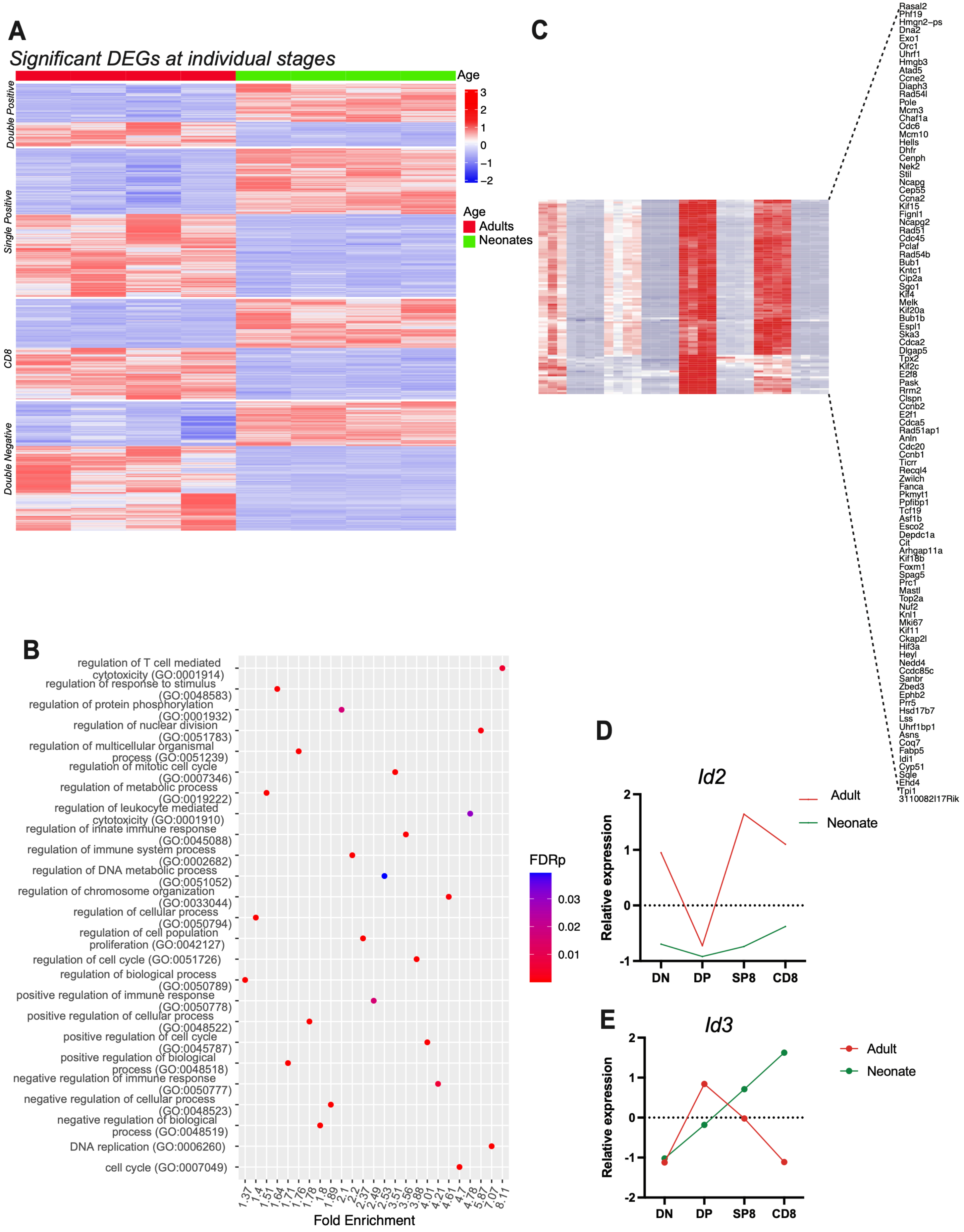

#### Figure S4

**A** *Significant DEGs across all developmental stages*

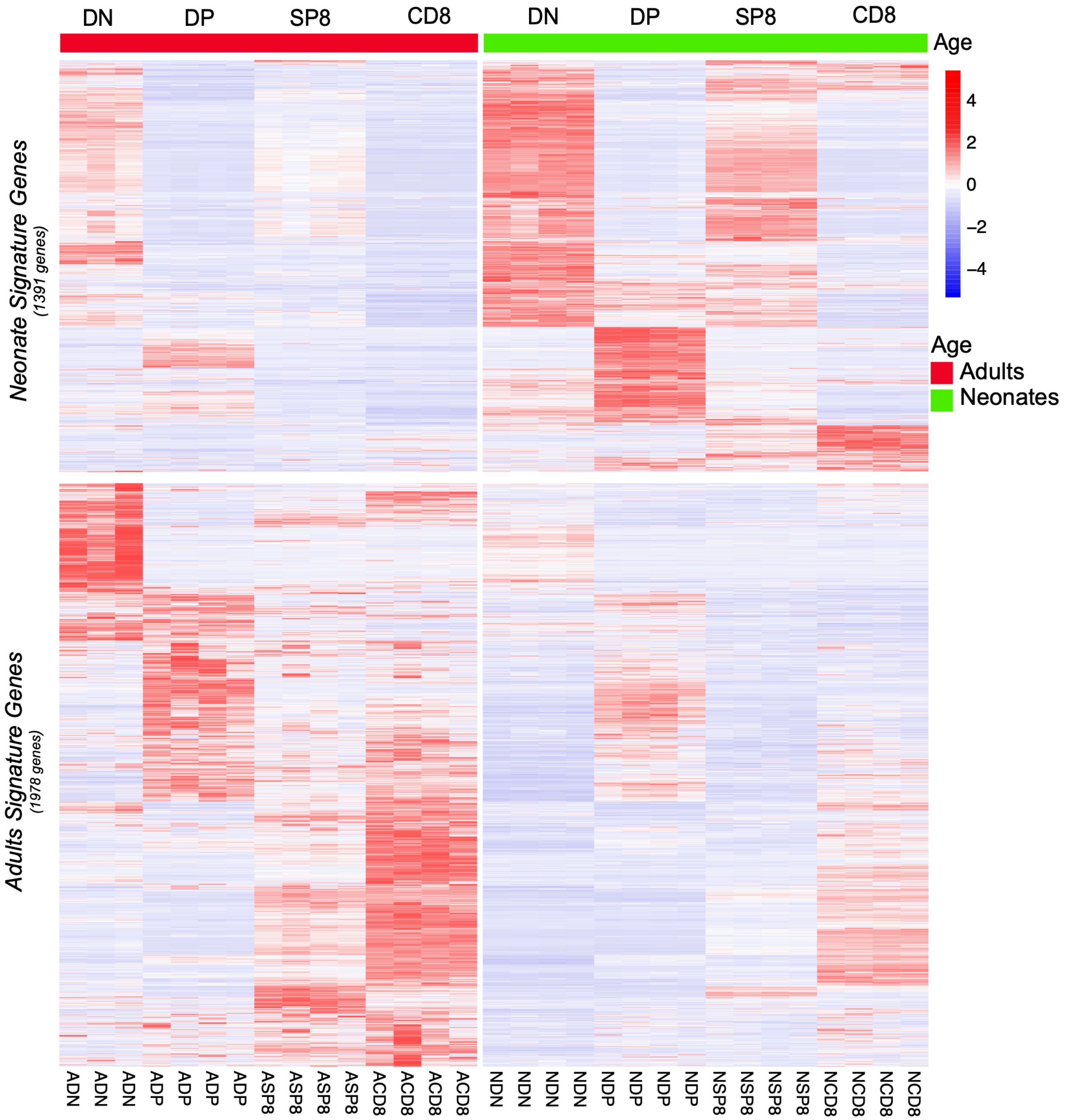

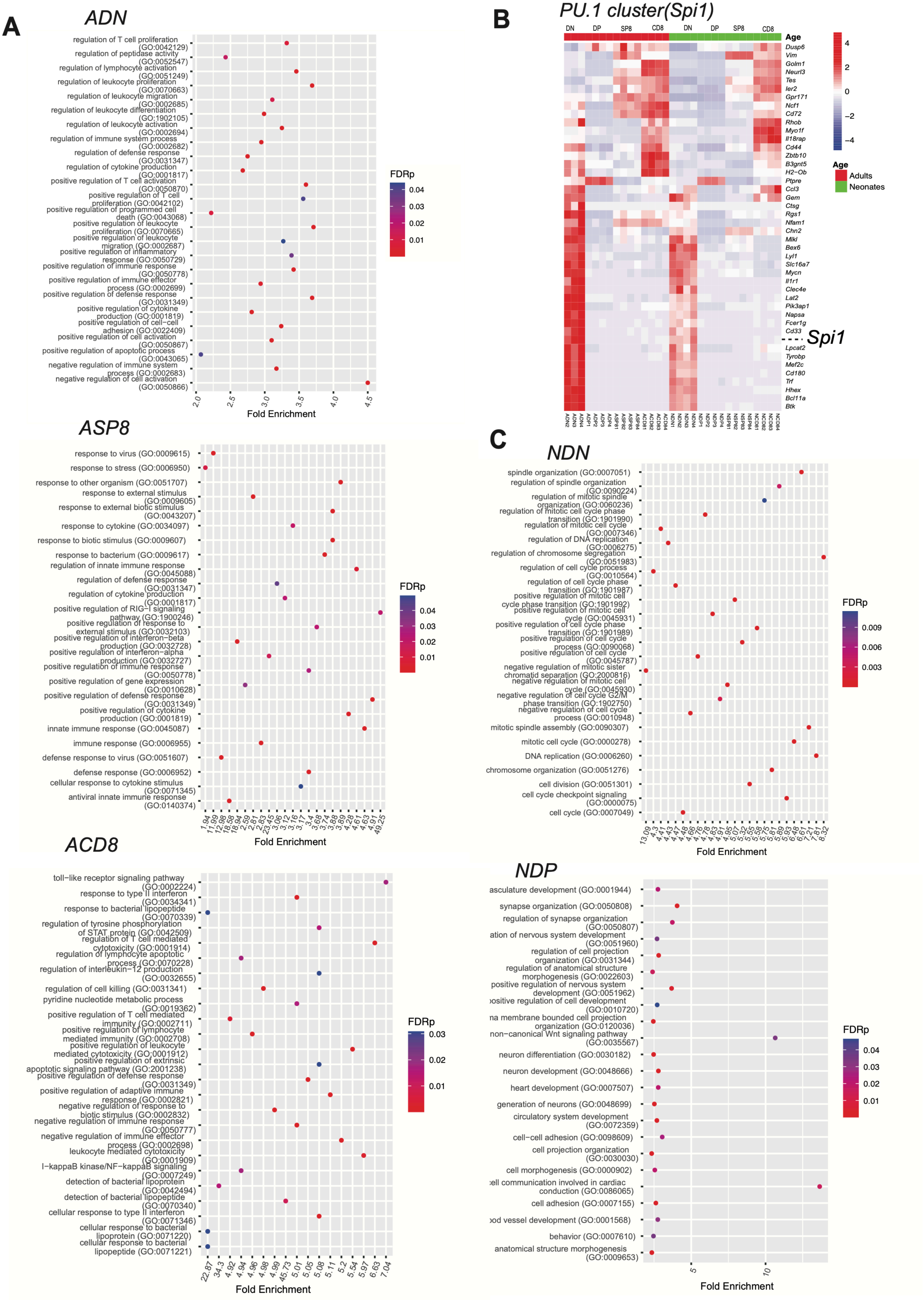

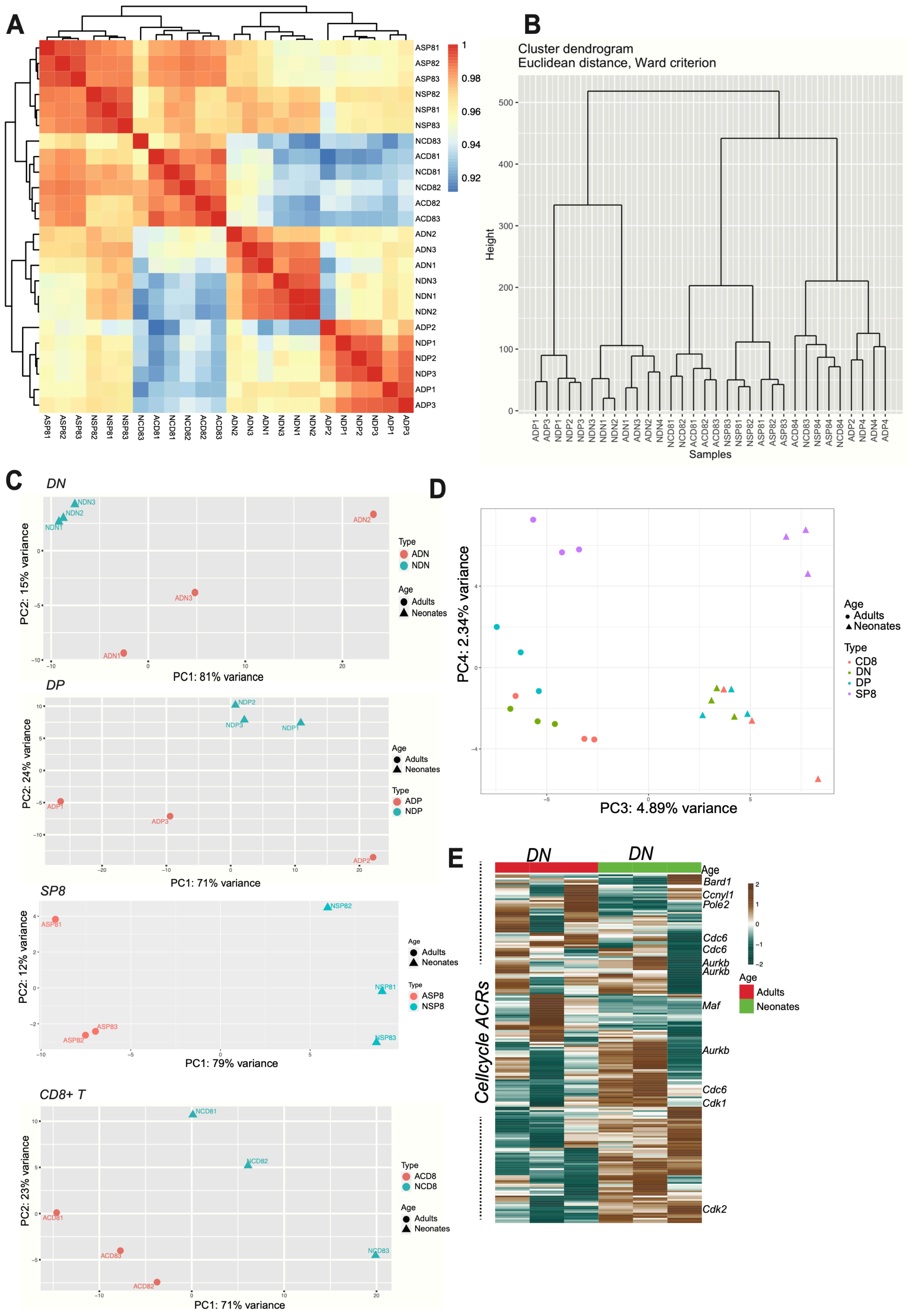

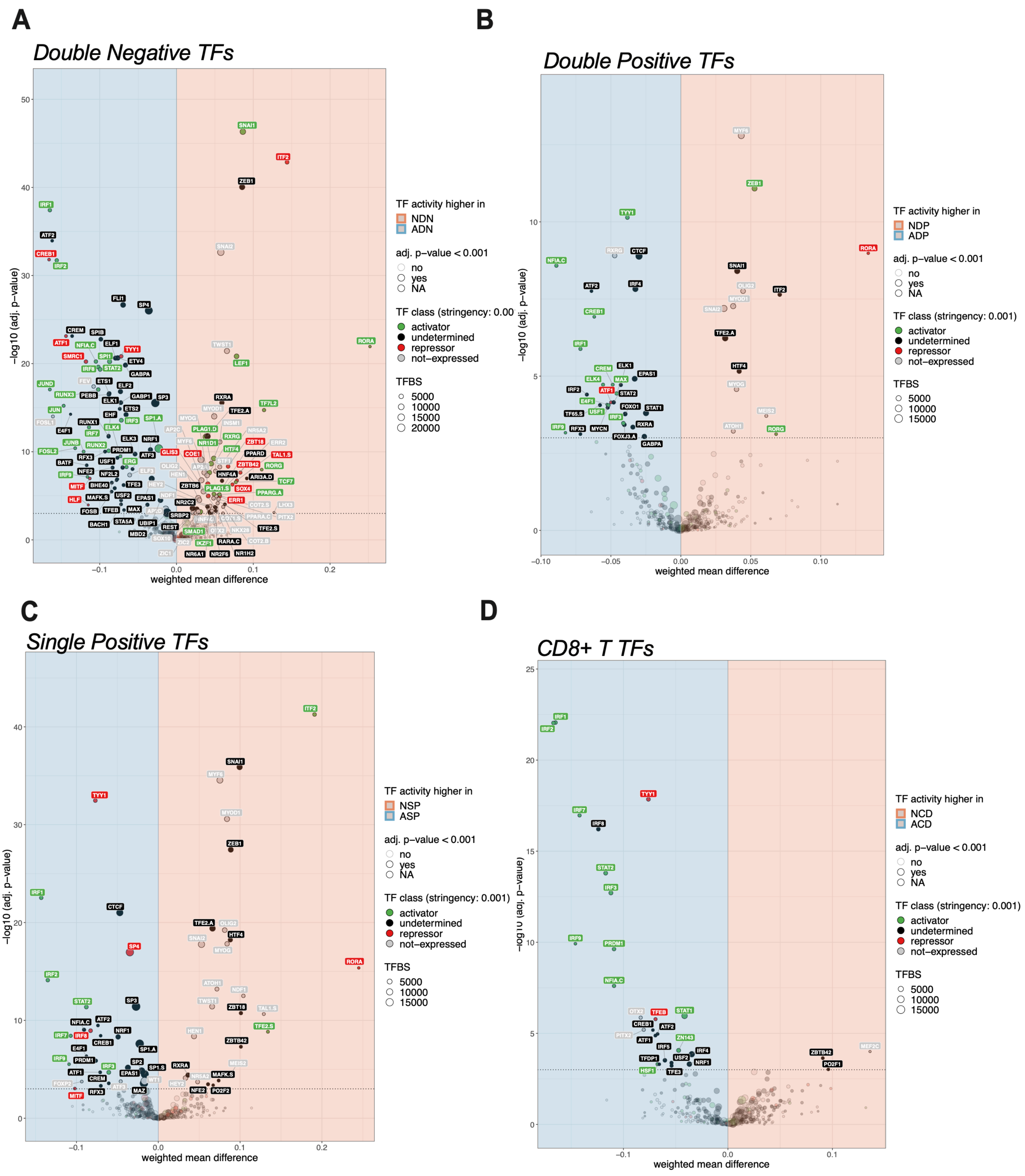

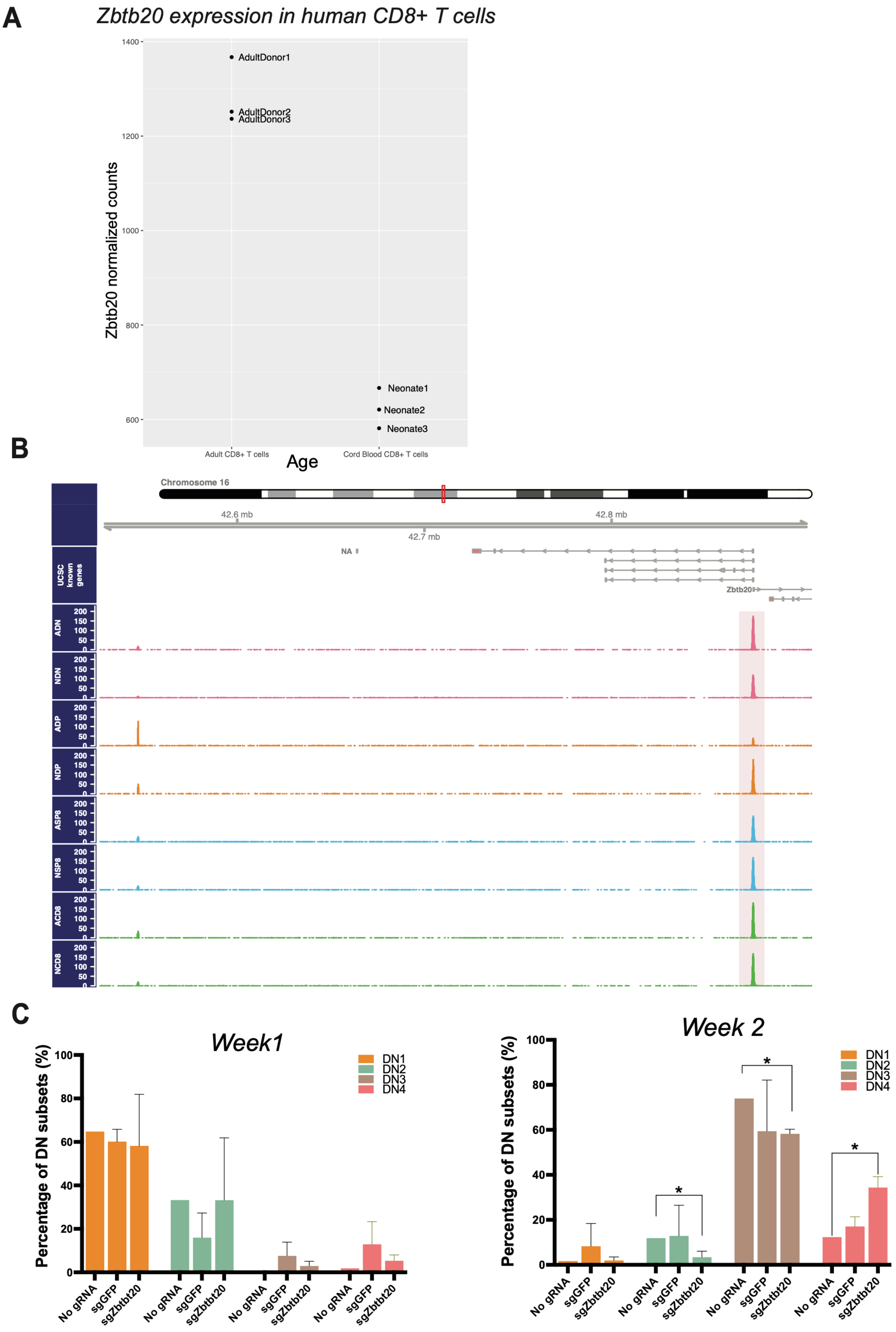

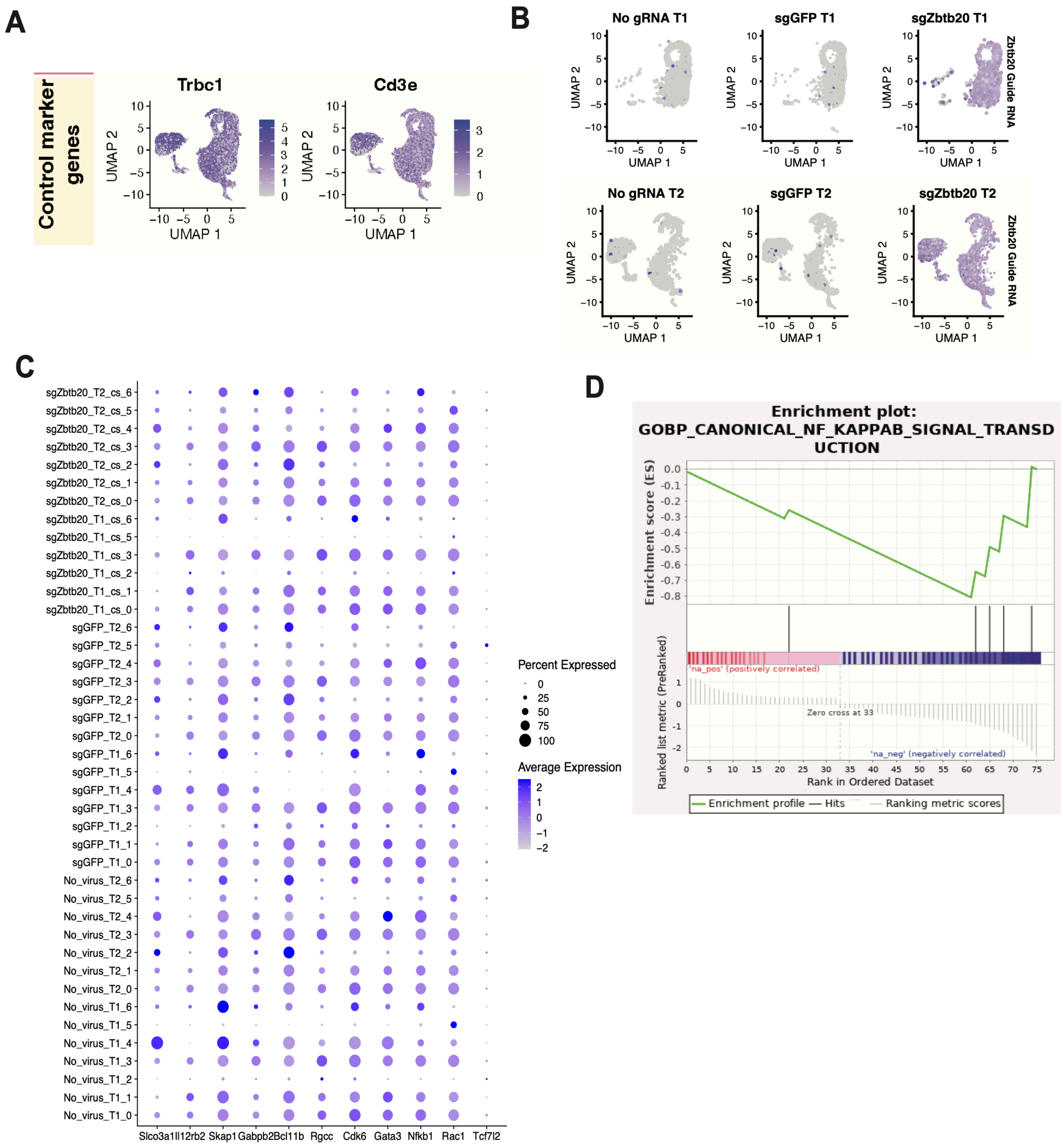
